# 5′ RNA Aminoacylation via Interstrand Acyl-Transfer

**DOI:** 10.64898/2025.12.18.695146

**Authors:** Josh Maiklem, James Attwater

## Abstract

The catalytic transformations driving coded protein synthesis revolve around linkage of amino acids to molecules of RNA as 2′/3′-aminoacyl esters. This defining molecular species of life is installed, supervised and utilised by complex catalysts, but its relative hydrolytic lability limits its accumulation and behaviour during the early development of the translation system in a simpler RNA-based biology. Herein, we describe a rapid catalytic transfer reaction inherent to RNA duplexes that establishes a more stable RNA-amino acid linkage. We observe spontaneous aminoacyl transfer from the 2′/3′-hydroxyl of a donor oligonucleotide to the 5′-hydroxyl of an adjacent template-bound acceptor oligonucleotide. In RNA systems dominated by base pairing, this transfer drives selective accumulation of the more stable 5′-aminoacyl RNA in the presence of a prebiotically-plausible aminoacylation reagent. This catalytic context could shape the distribution of aminoacylation amongst prebiotic RNAs and open opportunities for simple forms of RNA-directed peptide synthesis.

## Introduction

The central dogma of molecular biology is a cooperative endeavour between polymers of nucleic acid and peptide, but the origins of this arrangement remain unknown (1). In cells, the first step of biological ribosomal peptide synthesis is the charging of the 3′-terminus of tRNA with a cognate amino acid. The resulting tRNA 2′/3′-aminoacyl ester is the central actor in translation (2). This linkage a) facilitates the RNA-guided introduction of the amino acid into the ribosome peptidyl transferase centre (PTC), constituting the coded link between RNA sequence and amino acid identity; b) endows the amino acid with an RNA 3′-hydroxyl leaving group, activating it sufficiently for peptide bond formation whilst remaining resistant to uncontrolled polymerisation; and c) offers a vicinal 2′-hydroxyl, providing critical catalytic assistance in the ribosome PTC.

In biology, these processes are mediated by a series of complex, evolved catalysts. Aminoacyl tRNA synthetase enzymes (ARSs) first activate an amino acid as an aminoacyl adenylate using ATP, then transfer it onto the 2′- or 3′-hydroxyl of the tRNA to yield the terminal ester (2). Elongation factor Tu (EF-Tu) then recognises the aminoacylated tRNA, promotes the underlying isomerisation of the aminoacyl moiety from 2′ to 3′-hydroxyl (3), and (in complex with GTP) facilitates interaction with the ribosome-mRNA complex and decoding (4). The ribosome PTC first wields the aminoacyl ester as an amine nucleophile, and then in the peptidyl product as an ester electrophile, during iterative peptide bond formation (5).

In the face of this necessary catalytic complexity, it is unclear how coded peptides can be synthesised in simpler systems, in particular during the early evolution of the translation system itself. The informational and catalytic roles of RNA throughout biology’s translation system indicate that the molecular architecture of life arose out of an earlier biology run by RNA catalysts, culminating in the ribosome itself (6). However, little is understood about the nature of the transition from this ‘RNA world’ to an RNA-protein world. Proposed routes include stepwise emergence of function (7) via noncoded peptide synthesis (8) and/or co-option of existing ribozyme activity (9, 10) – but such models cite a pool of aminoacylated RNA ‘adaptors’ (e.g. tRNA) for use by a primitive peptidyl transferase ribozyme (1).

However, this stipulation is jeopardised by the hydrolytic lability of RNA 2′/3′-aminoacyl esters (11), particularly under the neutral/mildly basic conditions that promote aminoacylation and RNA polymerisation. In biology, aminoacyl-tRNA is protected from hydrolysis by EF-Tu, which excludes water from the vicinity of the ester bond whilst chaperoning it from its site of synthesis (ARS) to the ribosome where it will be used (12, 13). A primitive translation system could not have relied upon evolved macromolecules to protect prebiotically-aminoacylated RNA.

Nevertheless, RNA adaptors needed to maintain a high level of aminoacylation to prevent peptide product deletions or early termination and allow for the consistency of product needed to give proteocentric evolution a foothold. This problem would have been particularly acute given 1) the lack of any mechanism to gatekeep adaptor binding to mRNA based upon aminoacylation state (as implemented by EF-Tu (14)), and 2) the presumably slower nature of primitive peptide synthesis. Deacylated tRNA actively catalyses translation termination (10, 15), and even in *in vitro* ribosomal systems (with EF-Tu gatekeeping), translation is inefficient if a single tRNA exhibits below ∼20% aminoacylation (16).

While much research has focused on how 3′-aminoacyl oligonucleotides could have been generated prebiotically, less attention has been given to the stabilisation and retention of these as substrates for peptide synthesis. One potential route to this stabilisation is ‘chimeric’ aminoacyl-bridged RNA (17) formed via a phosphoramidate bond between the amine of a 3′-terminal aminoacyl oligonucleotide and the 5′-phosphate of an adjacent (template-bound) oligonucleotide, comprising an RNA duplex ‘nick’. This bridge substantially stabilises the aminoacyl ester (11) by avoiding the inductive effect of amine protonation under neutral/basic conditions. Such chimeric RNAs can form active ribozymes (18) and serve as a template upon which RNA can be synthesised by non-enzymatic polymerisation (19). However, by sequestering the amine moiety, this linkage precludes peptide synthesis without acid hydrolysis of the phosphoramidate to restore the (labile) 2′/3′-aminoacyl ester (20). The challenges associated with the biological form of aminoacylation have led to growing interest in a range of further substrates for RNA-guided prebiotic peptide bond formation, including mixed anhydrides, aminonitriles, thioacids, aminoacyl phosphoramidates, and ^6^A/^5^U-nucleobase-linked amides and carbamates (21–26).

Herein, we describe an RNA-mediated transesterification that rapidly converts 2′/3′-aminoacyl esters to a more stable form of aminoacylation – 5′-aminoacylation – that could have provided a prebiotic means of accumulating RNA-linked amino acids, whilst maintaining a generational relationship to extant 2′/3′-aminoacylation. Our observations suggest that this species is a direct outcome of prebiotic aminoacylation chemistry in base-paired RNA contexts. We explore the implications of this species’ formation and potential roles in models of the origins of translation.

## Results

### Routes to Stabilisation of RNA Aminoacylation

Modern biology relies upon evolved factors to stabilise and accumulate aminoacyl-tRNA. We set out to examine whether – in their absence – early molecular and environmental contexts could have achieved this. We first surveyed the half-life of a 2′/3′-aminoacylated RNA oligonucleotide (prepared using the flexizyme system (27), and fluorescently labelled at its 5′-terminus) under combinations of pH and temperature used to support ribozyme activity (Fig. 1*A* and Supporting Discussion, Fig. S1 and Fig. S2).

**Fig. 1.**
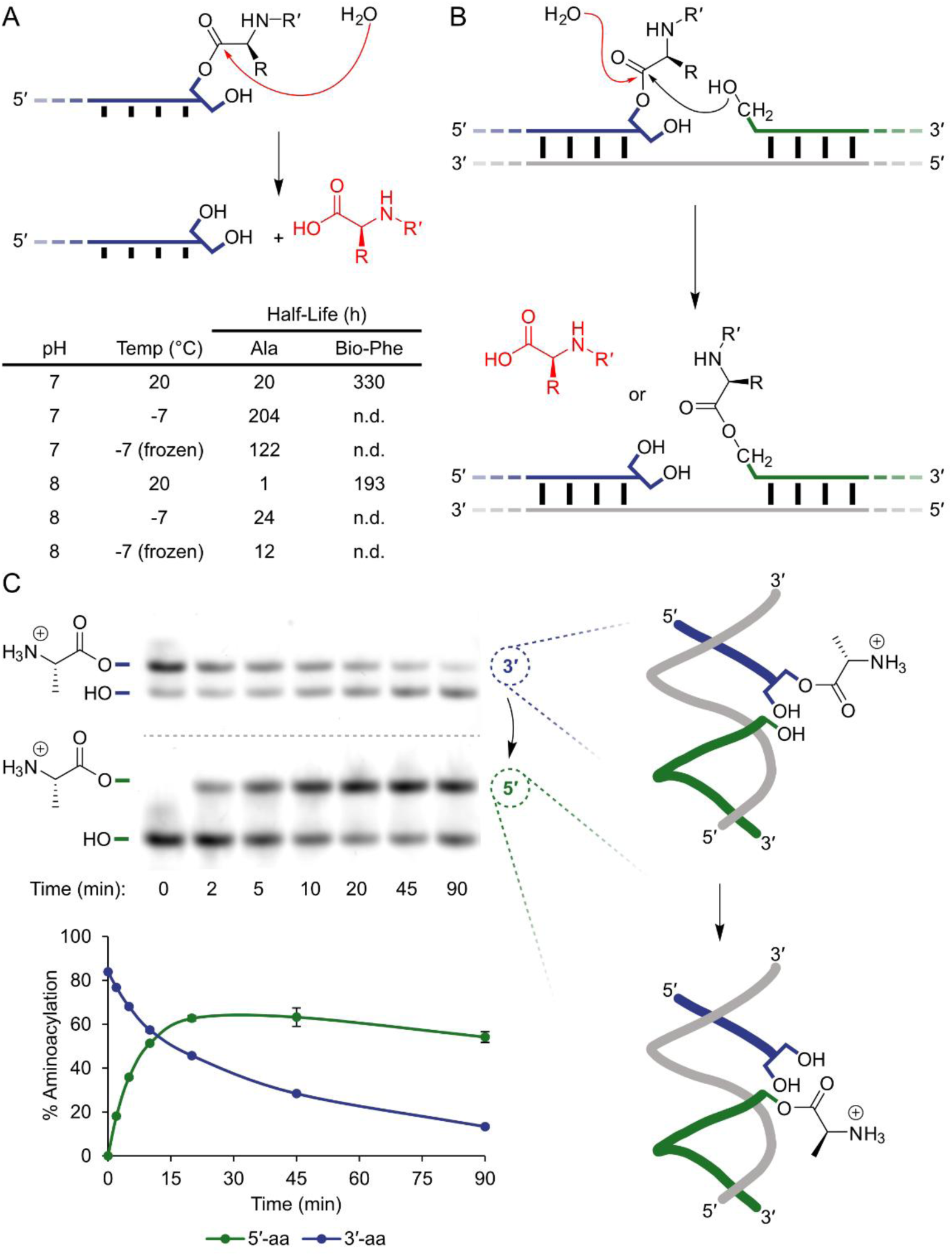
Behaviour of 2′/3′-aminoaycl RNA with or without an adjacent 5′-hydroxyl. (*A*) Direct hydrolysis of 2′/3′-aminoacyl RNA. Half-lives are shown for alanine (Ala; R=Me, R′=H) and N-biotinyl-phenylalanine (Bio-Phe; R=CH_2_Ph, R′=biotin) aminoacyl RNA (0.25 μM FITCPriCCA-2′/3′ ester, 50 mM 3-(N-morpholino)propanesulfonic acid (MOPS)·NaOH pH 7.0 or 8.0 (measured at 20°C), 200 mM NaCl, 0.05% Tween-20, 20°C or -7°C (supercooled or frozen)). Source data and quantification details in Fig. S1 and Supplementary Methods. (*B*) Schematic of the two reaction pathways available to a 2′/3′-aminoaycl ester located within a duplex nick. Hydrolysis (red arrow) leads to the free amino acid and an RNA oligomer with 2′/3′-diol. Aminoacyl transfer (black arrow) leads to 5′-aminoacyl RNA. (*C*) Transfer of alanyl ester from the 2′/3′-diol of an aminoacyl RNA to an adjacent 5′-hydroxyl across a duplex nick (pH 8, 20°C, 0.25 µM FITCPriCCA-Ala & TempA, 0.125 µM Cy5PriA). The 2′/3′-aminoacyl band of the FITC-labelled strand (blue) decreases with time (upper image) and a new 5′-aminoacyl Cy5-labelled strand (green) emerges (lower image). Observed %s of aminoacylation of each labelled RNA (n = 3, ± s.d.) sum to more than 100% in earlier timepoints due to the presence of excess FITCPriCCA-Ala. Right: 3-dimensional schematic of the nick construct showing the structure of the duplex facilitating aminoacyl transfer.

In particular, it was recently reported that frozen conditions lead to higher yields of ribozyme-catalysed and nonenzymatic RNA aminoacylation (24, 28). Freezing concentrates solutes into a supercooled eutectic phase, promoting RNA interactions and catalysis (29) and accelerating many prebiotically relevant reactions (24, 30–33). We established that freezing at -7°C increased alanyl RNA ester half-life to ∼5 days at pH 7 (Fig. 1*A*), a 6-fold stabilisation of the RNA aminoacyl ester compared to 20°C, and in line with more conservative estimates (34, 35) of aminoacyl ester temperature sensitivity. Interestingly, supercooling solutions to the same temperature was even more effective at stabilising aminoacyl RNA than freezing (Supporting Discussion).

More basic conditions (pH 8-8.5) support models of ribozyme-catalysed and non-enzymatic RNA polymerisation (36–39) (requiring 3′-hydroxyl deprotonation) and peptidyl transfer (involving aminoacyl amine deprotonation to form and productively resolve tetrahedral intermediate (40)). Here we observed a 10-20-fold decrease in the half-life of 3′-alanyl oligonucleotide vs pH ∼7 (Fig. 1*A*), consistent with a linear relationship between ester hydrolysis rate in ice and [^-^OH]. This stands in contrast to previous reports of aminoacyl nucleotide stability (11), which plateaued around the aminoacyl ester amine p*K*_aH_ (∼8 (41–43)) due to disproportionate acceleration of hydrolysis in its protonated form (34).

A peptidyl RNA mimic (the 2′/3′-ester of biotinyl-phenylalanine, in which amide *N*-capping precludes protonation (34)) showed surprising (>100-fold) stabilisation at pH 8 relative to phenylalanyl RNA (Fig. 1*A* and *SI Appendix* Fig. S1). However, the feasibility of peptide synthesis would be dominated by the stability of multiple *uncapped* aminoacyl RNA building blocks under conditions conducive to RNA catalysis (Fig. S3). Taken together, lowering temperature and pH can stabilise aminoacylation, but the increases in stability do not scale far beyond the known decreases in e.g. ribozyme reaction rates (25, 38). We therefore sought additional stabilising phenomena within the structure of RNA itself.

### Amino Acid Transesterification from 2′/3′- to 5′-Hydroxyls Across a Nicked RNA Duplex

Spontaneous hybridisation plays key roles in RNA replication, allowing copying of structured RNAs and RNA replication cycles, and may be reflected in the GC-rich nature of the postulated primitive genetic code (44). At low temperatures and raised solute concentrations (as found within the eutectic phase of ice) RNAs as short as trinucleotides predominantly exist in a hybridised state (45, 46), populating a variety of multistrand configurations involving gaps and nicks. The latter (Fig. 1*B*) arise from gapless hybridisation of oligonucleotides on a template strand and are further stabilised via coaxial stacking (stronger even than stacking within a contiguous helix (47)).

We tested the influence of this in-nick context upon the stability of 3′-terminal aminoacylation. Neighbouring 5′-phosphate and triphosphate groups had a small protective effect upon aminoacyl ester stability (Fig. S4). The most dramatic effect of this context, though, was not protective. When a dephosphorylated oligonucleotide – bearing a 5′-hydroxyl moiety – was sited next to the aminoacyl ester, we observed rapid and efficient deacylation in minutes.

To probe this phenomenon, we fluorescently labelled both the upstream (2′/3′-aminoacyl) and downstream (5′-hydroxyl) oligonucleotides comprising the nick. As the upstream oligonucleotide lost aminoacylation, we saw a new, matching downstream oligonucleotide band accumulate (Fig. 1*C*) in a 5′-hydroxyl- and template-dependent manner (Fig. S5*A*), indicating that the amino acid was in fact being transferred to the 5′-hydroxyl of the neighbouring strand via transesterification. This band persisted when all 2′/3′-aminoacylation was expected to have hydrolysed. Given its potential to prolong RNA-amino acid linkage, we investigated this transfer in more detail.

Aminoacyl transfer rates (k_tr_, judged from initial accumulation of 5′-aminoacylation) exceeded hydrolysis rates (k_hyd_) across different temperatures, with the k_tr_/k_hyd_ ratio being substantially higher at reduced temperatures (Fig. 2*A*). The transfer occurred cleanly even down to pH 6, >8 log units from the nucleophile’s p*K*_a_ (Fig. 2*B*), with the fastest transfer rates, as expected, being seen at pH 8 (6.64 h^-1^) and pH 9 (10.9 h^-1^) at 20°C. The transfer reaction is not dependent upon Mg^2+^ and was compatible with different buffers, buffer concentrations, and salts (Fig. S5*B*), implying that general base catalysis is not involved. The speed of the transfer is notable given the high p*K*_a_ (∼15) of the 5′-hydroxyl nucleophile.

**Fig. 2.**
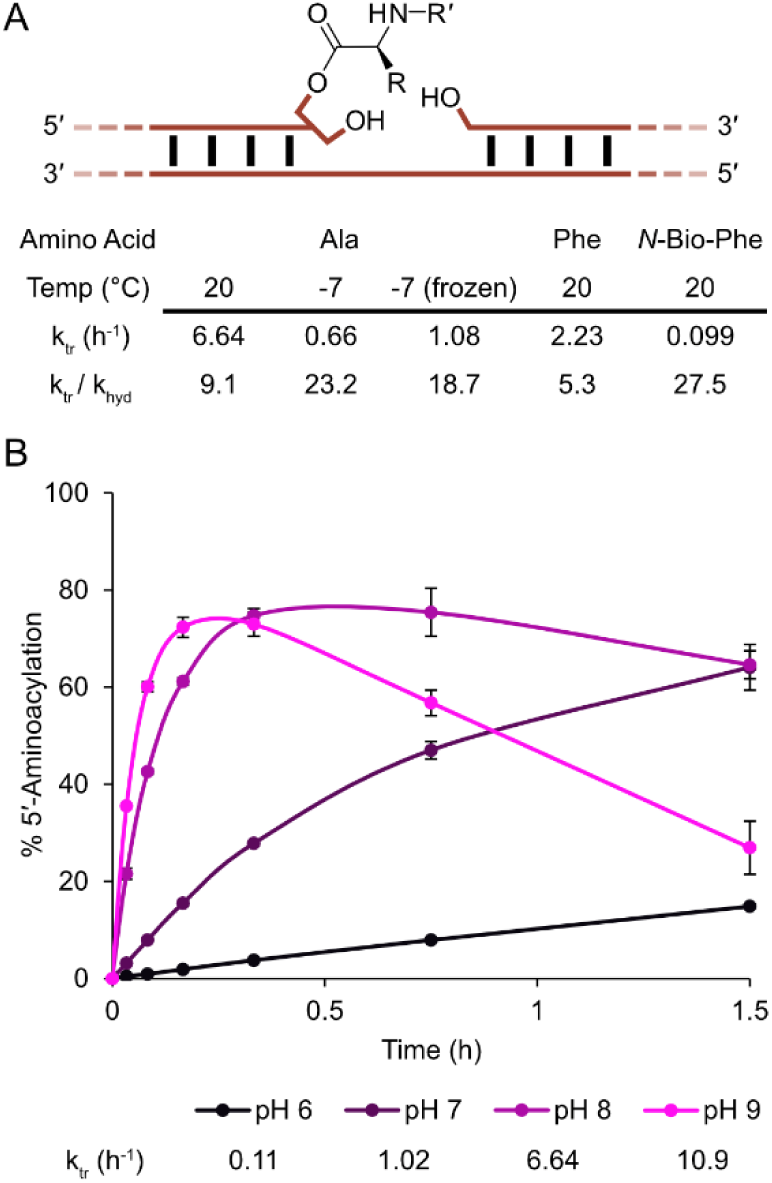
Kinetics of aminoacyl transfer. (*A*) Initial rates of transfer for alanine at different temperatures and phenylalanine and *N*-biotinyl-phenylalanine, all at pH 8.0, and comparison to the 2′/3′-aminoacyl ester hydrolysis rate in the same conditions. (*B*) pH dependence of aminoacyl transfer to the 5′-hydroxyl in a nick from 2′/3′-alanyl RNA (n = 3 ± s.d.). Initial rates are tabulated below. RNAs (0.25 µM FITCPriCCA-Ala & TempA, 0.125 µM Cy5PriA) were incubated at 20°C (200 mM NaCl, 0.05% Tween-20) in MOPS (50 mM, pH 7.0 or 8.0) or 2-(N-morpholino)ethanesulfonic acid (MES; 50 mM, pH 6.0) or N-cyclohexyl-2-aminoethanesulfonic acid (CHES; 50 mM, pH 9.0). Quantification details in *SI Appendix*.

In a nick, 2′/3′ → 5′ transfer dominated the fate of both alanine and phenylalanine, as well as the *N*-biotinylated derivative of phenylalanine (representing a peptide mimic) (Fig. 2*A*). The transfer rate was 3× faster for alanine than phenylalanine, mirroring the faster hydrolysis rate of alanine 2′/3′-esters (Fig. S1) (13, 35). A peptidyl moiety transferred much slower (23× for *N*-biotinyl-phenylalanine vs. phenylalanine), and at rates (0.099 h^-1^, pH 8, 20°C) similar to the transfer of *N*-biotinyl-methionine previously investigated (0.096 h^-1^, pH 7.3, 20°C) (48) as a starting point for ribozyme catalysis.

Having established efficient transfer from the 2′/3′-diol to 5′-hydroxyl, we considered the fate of the newly formed 5′-aminoacylation. Purified 5′-alanyl RNA transfer product exhibited a half-life of 9.3 h at pH 8, 20°C, ∼10× longer than the biological 2′/3′-aminoacylation (Fig. S6*A*), and a similar enhancement to that observed in an analogous, chemically synthesised 5′-phenylalanyl species (49). This stability likely reflects the higher p*K*_a_ of a 5′-hydroxyl leaving group (∼15) relative to that of the 2′/3′-diol (12-13 (50)) at the 3′-terminus, and the absence of a vicinal hydroxyl that could participate in hydrolysis.

Given the similar chemical natures of substrate and product, and potential operation of an equilibrium, we felt it prudent to investigate the possibility of the reverse (5′ → 3′) transfer, using purified 5′-aminoacyl RNA hybridised alongside an unmodified 2′/3′-cis-diol to act as the aminoacyl acceptor. However, we saw no evidence for the back-transfer. If any 2′/3′-aminoacylated RNA was generated, it either returned to the 5′ before it reached detectable levels, or hydrolysed. The latter pathway would represent 3′-terminus-mediated catalysis of 5′-aminoacyl ester hydrolysis. However, in these reactions we saw no change in the rate of loss of amino acid from the 5′ in the presence of an adjacent 3′-terminus, providing an upper limit on 5′ to 2′/3′ transfer of 0.0098 h^-1^ (Fig. S6*B*), <0.15% of the forward transfer – emphasising the relative strength of the 5′-ester linkage.

### RNA Context-Dependence of Aminoacyl Transfer

The robust nature of this transfer in competition with water likely reflects positioning of the 5′-hydroxyl nucleophile, orchestrated by RNA hybridisation, in which the tetrahedral intermediate of the transfer would bear steric resemblance to a standard RNA phosphodiester backbone linkage. Indeed, disruption to the geometry of the duplex by replacement of RNA template and/or 5′-acceptor strand with DNA versions dropped the transfer rate by up to 60-fold (Fig. 3*A*).

**Fig. 3.**
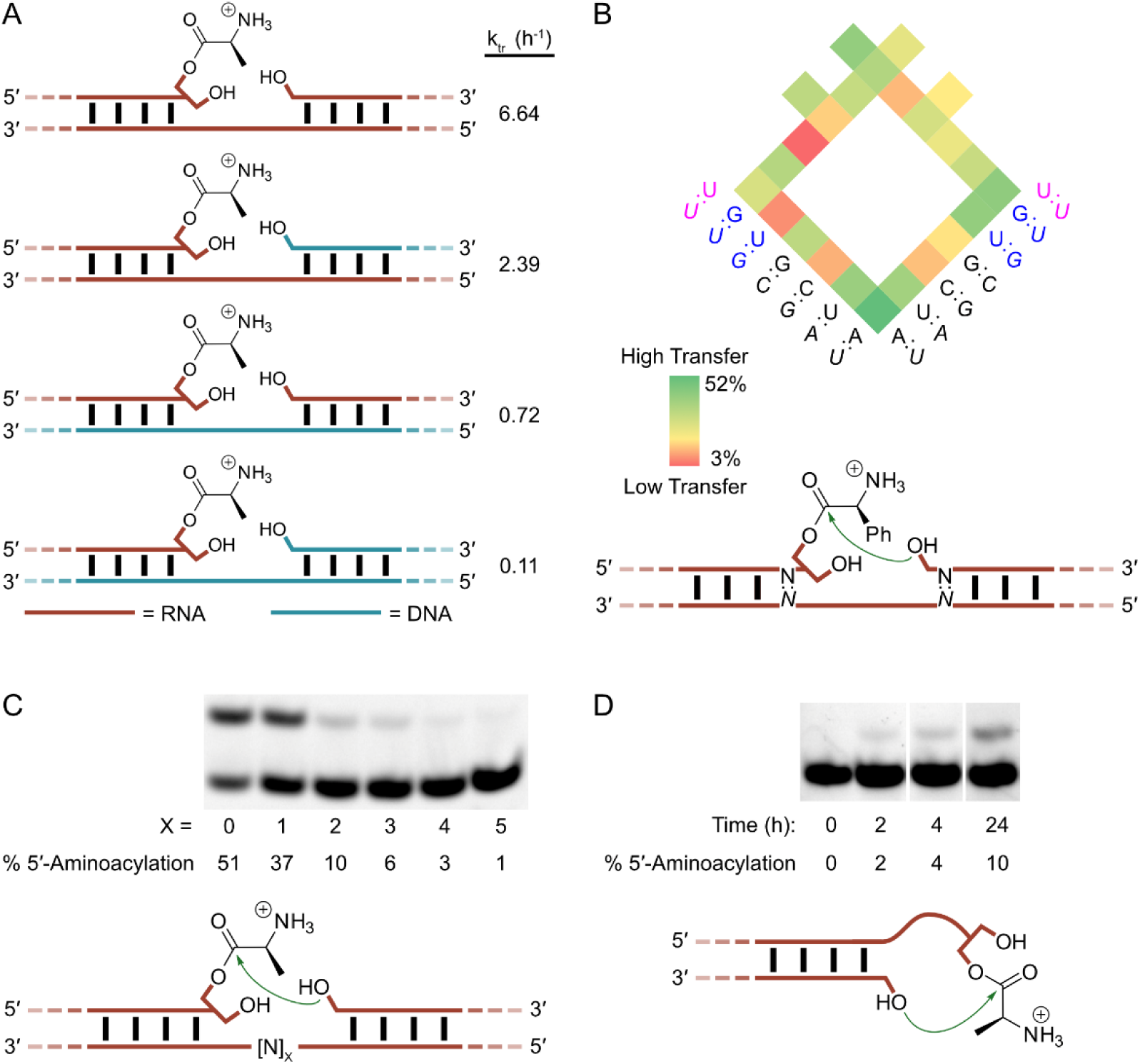
Interstrand aminoacyl transfer in different structural conditions. (*A*) Initial rates of transfer of alanine at pH 8, 20°C for a duplex consisting of all RNA oligonucleotides or having a DNA acceptor and/or template strand. (*B*) The dependence of transfer on sequence at the nick. Level of 5′-aminoacylation after 24 h at pH 8, -7°C (frozen) are shown for various combinations of bases at the four positions closest to the transfer site (for full data see Fig. S4). Watson-Crick base-paired sequences are shown in black, wobble base-pairs in blue and mismatches in pink. (*C*) 5′-aminoacyl transfer after 24 h at pH 8, -7°C (frozen) across gaps of varying length. Templates contained an intervening stretch of random sequence ([N]_x_ where x=0-5) between the donor and acceptor hybridisation sites. (*D*) Aminoacyl transfer occurs to a paired 5′-hydroxyl from a 2′/3′-aminoacylated GCCA-3′ overhang, pH 8, -7°C (frozen).

Within the RNA context, we sought to investigate the influence upon transfer of nucleotide identity flanking the nick that might exert more subtle effects upon double helix structure and 5′-hydroxyl nucleophile positioning (51). We found that the flexizyme system could be used to synthesise 2′/3′-phenylalanyl oligonucleotides terminating in each of the four canonical ribonucleotides (Fig. S7), allowing us to test the transfer rates from the four 3′-terminal bases to downstream oligonucleotides with varying 5′-residues. These oligos were hybridised to perfectly complementary template strands, as well as strands inducing a range of mismatches (Fig. 3*B* and Fig. S8). All other aspects of the system were kept constant to ensure comparability.

After 24 h incubation, we observed a range of transfer extents across the different junction sequences. The most extensive transfer appeared to be 3′-A:U → 5′-A:U (52% 5′-aminoacylation). Nicks bordered by canonical Watson-Crick base-paired residues exhibited median 33% transfer product, and a transfer to total hydrolysis ratio of 0.7, but multiple combinations transferred at this level, including wobbles on either or both sides of the nick, as well as mismatches. Interestingly, a 3′-A:U → 5′-G:U wobble transfer proceeded better than a 3′-A:U → 5′-G:C transfer (42% vs 18%); on the other hand, the least efficient transfer was 3′-G:U → 5′-C:G (3%). The sensitivity of transfer rates to small changes in sequence, but with little apparent correlation to base identity either side of the junction, implies a critical role for positioning of the 5′-hydroxyl in the transfer, with some contexts seemingly limiting its access to the aminoacyl ester.

In a ‘prebiotic’ oligonucleotide mixture, correctly base-paired junctions would likely have coexisted with a range of other junction architectures including loops and overhangs. Therefore, we explored the generality of the 2′/3′- → 5′-aminoacyl transfer in these divergent contexts. We first tested aminoacyl transfer across a template gap of various lengths in an RNA complex resembling those above but using RNA templates with intervening random sequence ([N]_x_, x=0-5) between donor and acceptor oligonucleotides (Fig. 3*C*). The transfer reaction diminished as the two reactive termini were moved apart, but significant transfer persisted across a gap of 1 nucleotide. This persistence of activity concords with the observation that template-bound oligonucleotides interact across 1-nt (but not 2-nt) gaps in ways that stabilise their hybridisation (47).

Recent work has demonstrated the potential for specific overhang sequences to facilitate reactions between RNA termini, perhaps by assumption of discrete structures (52–54). We investigated transfer at a version of the microhelix, a tRNA acceptor stem mimic. Transfers using more reactive substrates have been observed within this context, including aminoacyl transfer from a 5′-phospho-carboxy mixed anhydride to an overhanging 3′-hydroxyl (55), or to the amine of a 2′/3′-aminoacyl ester (23). We observe slow but distinct aminoacyl transfer to 5′-OH in this system, with a GCCA-3′ overhang (Fig. 3*D*), confirming the potential for the loop transfer phenomenon to deacylated such overhangs. However, activity was abolished by a single-nucleotide substitution in the overhanging sequence (to ACCA-3′).

### Modelling of Prebiotic Aminoacylation and Transfer

We returned to the more abundant nick context to understand the implications of transfer for aminoacylation in prebiotic systems. Across tested conditions and pHs, the rate constants for the hydrolysis or transfer of amino acids within the 3′-A → 5′-A junction suggest that 2′/3′-aminoacylation partitions to the 5′-hydroxyl 5-28x faster than it hydrolyses. We integrated the set of measured constants (pH 8, 20°C) into a simple model that recapitulates the accumulation of 5′-aminoacylation observed in our experimental data (Fig. S9) and predicts a maximum 5′-aminoacylation of 86% after 36 mins, with 58% remaining after 6 hours, starting from fully 2′/3′-aminoacylated RNA (Fig. 4*A*). Without the transfer parameter, only 1% 2′/3′-aminoacylation remained after this time. The transfer allows RNA to capture the amino acid in a hydrolysis-resistant state.

**Fig. 4.**
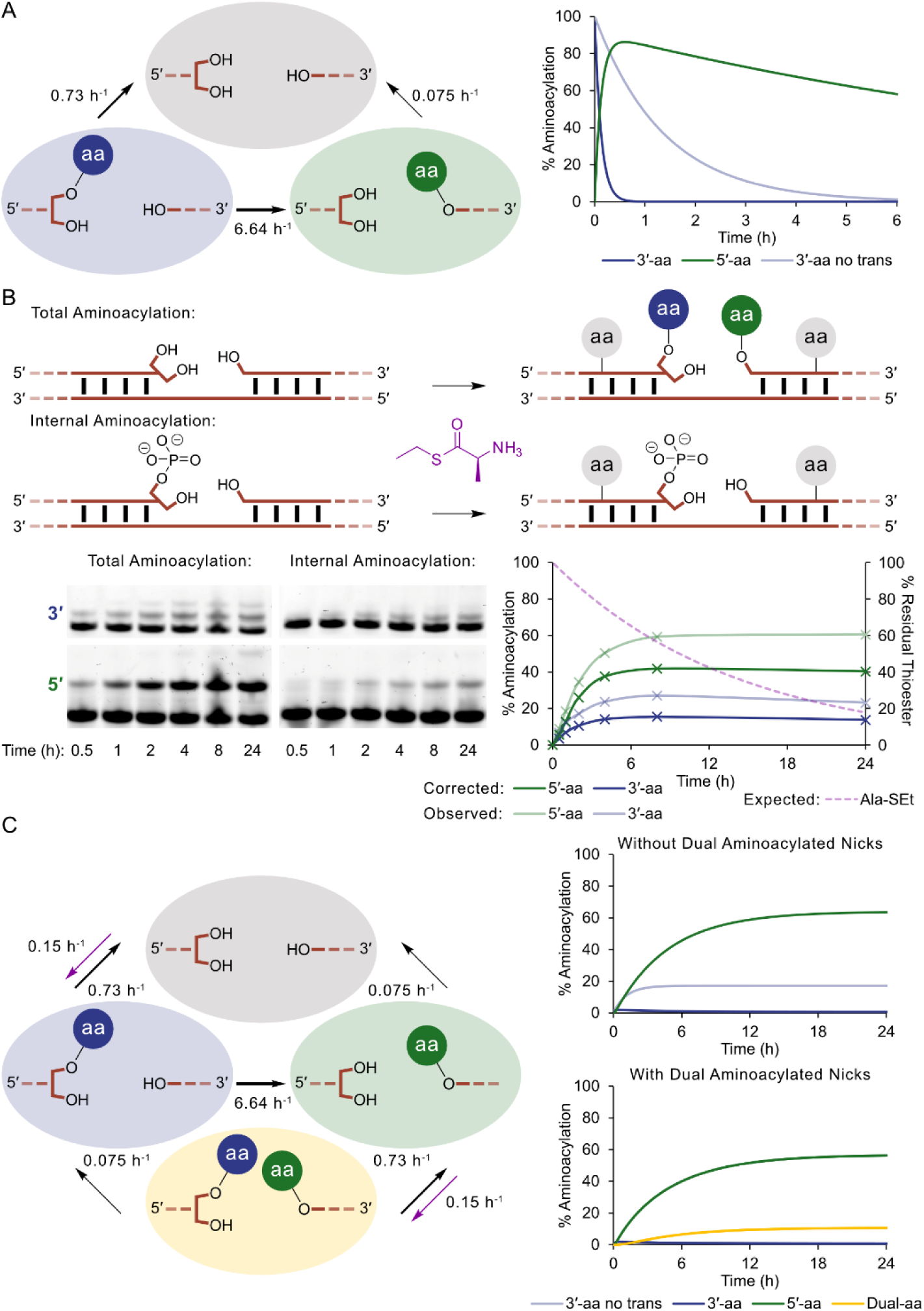
Modelling of nick system behaviour with and without constant aminoacylation level. All models use rates calculated at 20°C pH 8 (200 mM MOPS, 200 mM NaCl, 0.05% Tween 20). Thioester aminoacylation rates were measured in the same conditions with L-Ala-SEt (200 mM). Rates are stated next to arrows as pseudo first order observed rates. (*A*) Simple model of a junction system, with no constant aminoacylation, starting with 100% of nicks 2′/3′-aminoacylated, plotted against 2′/3′-aminoacylation for model system without transfer (light blue). (*B*) Above: oligonucleotide systems for measuring total (in-nick and internal 2′) aminoacylation, starting with no aminoacylation but 200 mM L-Ala-SEt thioester. Thioester does not aminoacylate the terminus of 3′-phosphoryl RNA (24). Below left: thioester aminoacylation in these systems (0.25 μM RNAs, 10% excess of template). Below right: levels of observed total aminoacylation of each oligonucleotide and ‘corrected’ terminal aminoacylation in the nick (by subtraction of observed internal aminoacylation). Estimated residual thioester % is also shown in purple. (*C*) Left: junction model incorporating transfer and dual-aminoacylation of junctions. Right: predicted accumulation of terminal aminoacylation by thioesters assuming a constant 2′/3′- aminoacylation rate of 0.15 h^-1^. The graphs show that higher levels of steady-state 2′/3′-aminoacylation are expected only when the dual-aminoacylated nick state is included in (below) vs excluded (above) from the model.

A fully 2′/3′-aminoacylated starting state is unrealistic, given the challenging nature of abiotic RNA aminoacylation. To properly understand the impact of this transfer in prebiotic scenarios, we needed to model steady state RNA aminoacylation, incorporating an empirical aminoacylation ‘on’ rate.

Ribozyme-catalysed 2′/3′-terminal aminoacylation (for example by the flexizyme system) can achieve high yields of aminoacyl-RNA (28), but faces challenges reconciling amino acid generality and enzyme:substrate turnover. The interplay between turnover and abundance of these evolved catalysts renders RNA-catalysed aminoacylation challenging to model in a steady state.

A nonenzymatic aminoacylation route using activated amino acid appeared more tractable, and more applicable to pools and complexes of mixed-sequence RNAs serving as an early context for the emergence of translation. However, the use of biological activated amino acids - mixed anhydrides – is not necessarily effective, as these tend to react better with CO_2_, amines, water, or other species in the molecular milieu (56).

Recently, reports of microhelix aminoacylation by cyanomethyl ester-activated amino acids (33) and phenol esters (16), and efficient, prebiotically plausible nucleoside and oligonucleotide 2′/3′-aminoacylation by thioester-activated amino acids (24) have reopened this possibility. In particular, thioesters exhibit chemoselectivity towards aminoacylation at RNA hydroxyls over competing amines. They preferentially aminoacylate 2′- and 3′- over 5′-hydroxyls, and in an RNA duplex target terminal 2′/3′-hydroxyls over internal 2′-hydroxyls (24). At pH 8 such thioesters drove ∼10% 2′/3′-aminoacylation/hour, and at pH 6.5 (where product aminoacyl RNA is more stable) could establish up to ∼50% nucleoside aminoacylation in the eutectic phase.

To assess compatibility of thioester-driven aminoacylation chemistry with aminoacyl transfer, we incubated 200 mM L-alanine ethanethiol thioester (Ala-SEt) with the unmodified dual-labelled RNA nick construct (3′-A-OH, 5′-A-OH, 20°C). All RNA oligonucleotides therein were fully base-paired to attenuate potential off-target aminoacylation at internal 2′-hydroxyls (24). We observed effective aminoacylation from thioesters, but selectively accumulating on the 5′-hydroxyl alongside minor levels of 2′/3′- and internal 2′-aminoacylation (Fig. 4*B*).

The thioester chemistry is unable to directly yield stable 5′-aminoacylation at appreciable levels (24). In the absence of the junction, incubation with Ala-SEt of the individual labelled oligonucleotides (though paired to complementary RNAs) resulted in a (initially) higher level of 2′/3′-aminoacylation but no 5′-aminoacylation (Fig. S10). The pseudo-intramolecular context of the transfer reaction is critical for 5′-aminoacylation under these conditions.

This aminoacylation reaction does not fully constitute a steady state due to slow (t_1/2_ ≈ 10 h, pH 8, room temperature (24)) depletion of the thioester substrate by hydrolysis, limiting 5′-aminoacylation to 40%. We used a version of this system where no transfer was possible (possessing a 5′-phosphate adjacent to the 2′/3′-diol; Fig. S11) (24) to measure the initial rate of 2′/3′-aminoacylation by thioester (pseudo-first order rate constant: k_obs_ = 0.15 h^-1^, 200 mM thioester pH 8, 20°C). Using this, we modelled the effect of fixing this concentration of activated thioester. Starting from unacylated termini, this predicted a steady state aminoacylation level of 64% 5′- and 1% 2′/3′-, reached after around a day (Fig. 4*C*). In the absence of transfer, only 17% 2′/3′-aminoacylation is maintained; 64% 2′/3′-aminoacylation would require a prohibitive 1.7 M sustained concentration of thioester (Fig. S11).

### Colocalisation of Amino Acids in a Junction

Interestingly, we observed a much higher level of 2′/3′-aminoacylation than this initial model predicts. A slightly lower level would be expected in the model as it does not account for the 10% experimental excess of template (slightly lowering 2′/3′-aminoacyl loss). But neither accounting for this, nor integrating a back-transfer term (even at 0.1 h^-1^, >10x higher than the calculated maximum (Fig. S6*B*)) into the model explained the observed 2′/3′-aminoacylation.

However, including continued 2′/3′-aminoacylation in 5′-aminoacylated junctions better matched the observed levels (Fig. 4*B* and *C*). This implies that after 5′-transfer, thioester re-aminoacylates 2′/3′-hydroxyl termini within the same nick, generating a population of dual-aminoacylated RNA junctions. This increases model aminoacyl loading to 0.79 per junction, and yields a population of ∼11% junctions with adjacent amino acids.

Iterative peptidyl transfer has not been observed in the absence of the ribosome – partly because of the intrinsic catalytic challenge of this reaction (34), and partly because of the apparent absence of accessible structural contexts in which two RNA aminoacyl esters are naturally juxtaposed. Bis-2′/3′-aminoacylated tRNA termini have been observed in biology (57), but these show no evidence of intramolecular peptide bond formation. Our observation of junction dual-aminoacylation via aminoacylation/transfer/reacylation led us to consider whether the RNA-mediated colocalisation established by the junction context could promote peptide bond formation. In particular, a dual-aminoacylated junction possesses similar nucleophile/electrophile spacing and geometry to the previously reported formation of amino acid-bridged chimeric RNA molecules by RNA-templated attack of a 2′/3′-aminoacyl amine upon a reactive 5′-phosphorimidazolide (17, 53).

Should such a context prove advantageous for peptide bond formation, in the absence of catalytic assistance, it would still likely be inefficient due to the inherent difficulty in outcompeting hydrolysis (34). However, if a ribozyme could take advantage of the proximity effects brought about by a dual-aminoacylated nick, such a catalyst may be able to access cycles of iterative peptide growth in concert with spontaneous aminoacylation and 3′ to 5′ transfer reactions (Fig. 5).

**Fig. 5.**
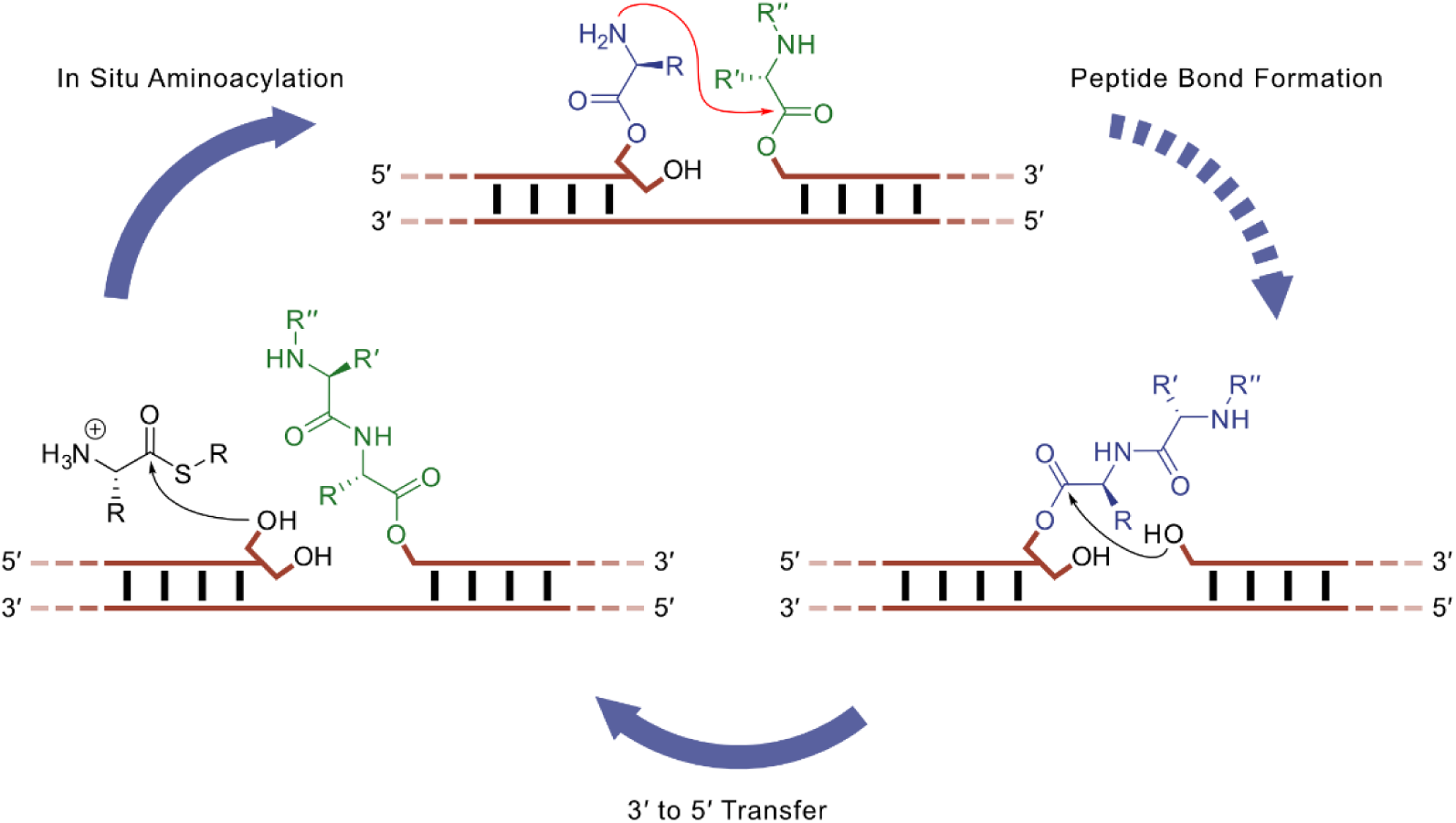
A putative iterative peptidyl synthesis cycle supported via thioester/ester transfer chemistry. In situ aminoacylation and 3′ to 5′ transfer are spontaneous; catalysis of peptide bond formation would restart the cycle. The multistep conversion of thioester to amide would drive the cycle energetically, under the control of RNA through the selectivity of thioester reaction with RNA 2’,3’-hydroxyls, the transfer to the 5′, and nick-mediated positioning and catalysis of peptide bond formation.

## Discussion

The instability of the biological forms of RNA aminoacylation, the 5′-phospho-carboxy mixed anhydride and 2′/3′-aminoacyl ester, jeopardises their use in ancestral forms of translation (58). Here we show that a more stable form – the 5′-aminoacyl ester – is readily accessible via a template-catalysed transfer from the more labile 2′/3′-aminoacyl ester. Indeed, under conditions of direct chemical aminoacylation, 5′-aminoacyl ester formation is the predisposed outcome (∼6:1) in junctions of base-paired RNAs. This broadens opportunities for primordial RNA systems to accumulate and wield aminoacyl esters, including for peptide bond formation itself.

Unlike other alternative prebiotic RNA-amino acid linkage chemistries, 5′-aminoacylation retains a linkage to biological 2′/3′-aminoacylation through its obligate intermediacy via the transfer reaction. This provides a rationale for a system reliant upon 5′-aminoacylation to transition to the modern chemistry by short-circuiting this pathway and exclusively using 2′/3′-aminoacyl esters for peptide bond formation, once it has developed catalysts to synthesise and manipulate such RNAs.

Indeed, given its hydrolytic instability, life’s choice of aminoacyl RNA linkage chemistry appears puzzling. It can be stabilised by low temperatures, but is substantially destabilised by the higher pHs employed in models of template- and ribozyme-catalysed RNA polymerisation, and that accelerate its formation from activated amino acids – constraining the level of 2′/3′-aminoacylation achievable before enzymes.

This challenge is amplified when considering RNA template-bound adaptors for iterative peptide synthesis. In a toy model of translation from contiguous template-bound RNA adaptor molecules (Fig. S3), a 2′/3′-aminoacylation rate of 0.15 h^-1^ and hydrolysis rate of 0.73 h^-1^ (the experimentally determined rates for alanine at pH 8, 20°C) establishes a steady state level of 17% 2′/3′-aminoacylation per adaptor. Yet this corresponds to only 2×10^-6^% concurrent aminoacylation of all ten template-bound RNAs needed for successful peptide synthesis. Applying our kinetic model of junction aminoacylation, including transfer to the 5′, a steady state level of 67% 5′-aminoacylation per adaptor is achieved. This corresponds to 1.8% of templates in an analogous system exhibiting the full aminoacylation across all adaptors, an increase of six orders of magnitude. An aminoacylation rate of ∼1 h^-1^ would yield 56% of such templates bound by fully-aminoacylated adaptors.

Oligonucleotide 5′-phosphate species – key building blocks for nucleic acid replication – do not form junctions competent for this transfer. However, given the challenges of prebiotic nucleoside/nucleic acid 5′-phosphorylation (59, 60), and the fact that 5′-hydroxyls are also products of RNA hydrolysis, 5′-hydroxyl groups are likely to have been prevalent in the prebiotic milieu. Indeed, RNA ligation may have first proceeded through the attack of 5′-hydroxy-terminal RNA into 2′,3′-cyclic phosphates (61). In environments where oligonucleotides predominantly adopt base-paired states (45, 46), nicks harbouring abutting 5′-OH and 2′/3′-OH termini would be frequent and 5′-aminoacylation emerges from aminoacylation chemistry.

Despite different rates of transfer of different amino acids, we observed very similar rates of L-alanine and D-alanine transfer (Fig. S12). The RNA duplex provides a chiral environment that has been reported to confer a degree of stereoselectivity upon some reactions (20, 55, 62, 63), but the unphosphorylated junction site used here would be expected to be relatively solvent-exposed. The transfer reaction is, though, quite sensitive to duplex structure, with a decreased rate of transfer on a DNA template or to a DNA 5′-hydroxyl (despite the similarity of the 5′-terminus of DNA to that of RNA). This increases the scope of the transfer under potential concurrent prebiotic presence of DNA with RNA (64).

The transfer efficiency was notably influenced by the combinations of the nucleobases flanking the nick. Though the transfer performs best with flanking adenosines, G or U residues on each side are found in both low- and high-efficiency transfer nicks. We hypothesise that this phenomenon results from specific stacking interactions across the nick. In particular, the observed extent of transfer across each junction sequence is highly correlated (Fig. S13) with the free energy change of +1 lengthening of a helix via that sequence (used in nearest neighbour predictions of RNA folding (65)) as well as (to a lesser extent) the measured stabilisation of corresponding coaxial stacking upon hybridisation (47). Unexpectedly, these correlations were negative: the more stabilising the effect of helix propagation (from stacking and hydrogen bonding), the less transfer occurred in the equivalent nick.

As stacking energies of nucleobases in a nick are greater than in contiguous helices and are realised at warm temperatures (47), we expect that all such junctions exist in a predominantly stacked state in our assays. Potentially, the specific base stacking conformation mimicking that in an intact duplex leads to unfavourable positioning for this transfer, despite the resemblance of a phosphodiester’s geometry to the tetrahedral intermediate of the transfer. Nick nucleobase combinations accessing more favourable stacking in a coaxial vs helical context likely induce a different 5′:2′/3′-hydroxyl relative positioning that facilitates aminoacyl transfer. The variable rates of the transfer reaction emphasise the potential importance of junction sequence in other catalytic contexts including nonenzymatic ligation for assembly/replication of RNA.

During the screening of junction sequences, we found that flexizyme activity tolerated RNA substrates with a terminal ACCN sequence, accommodating all four terminal ribonucleotides (similar activity has been reported for r24, an ancestor of the flexizymes (66)). We attribute this to their mode of substrate recognition: flexizymes do not form a Watson-Crick base pair with the very terminal nucleotide, only the preceding three (67). Despite large differences in yields for both dFx and eFx with these substrates, the catalyst was sufficiently effective to allow preparation of phenylalanyl derivatives of all four termini (Fig. S8). Though this catalyst is known to aminoacylate the 3′-OH (68), aminoacyl exchange occurs rapidly between 2′- and 3′-OH with respect to hydrolysis (34, 69). It is therefore unclear whether the transfer exhibits a preference for one of these hydroxyls over the other.

We did not see evidence of the reverse (5′ → 2′/3′) transfer. The higher p*K*_a_ of the 5′-OH likely strengthens this ester bond, reflected in its slower hydrolysis, shifting any potential equilibrium between the two esters decisively towards the 5′. Interestingly, there is evidence that the transfer of a peptidyl ester in the ‘backwards’ (5′->2′/3′) direction can be catalysed by ribozymes. The ATRib ribozyme can catalyse partial equilibration of a peptidyl ester between the 3′-termini of two RNAs via its 5′-ester as an intermediate in a nick context (70). However, very low yields were obtained for peptidyl transfer to a nick 2′/3′-terminus from 5′-peptidylated ATRib (generated by an appended glutamine recognition domain using cyanomethyl ester-activated biotinyl glutamine) (70), reflecting the challenge for catalysts to change the position vs the rate of equilibrium.

Direct aminoacylation via free activated amino acids represents an intriguing alternative to achieving pool RNA aminoacylation. These would bypass the need for development of synthetase catalysts before the emergence of peptide synthesis that could harness them (71) (though other roles of aminoacyl-RNA, such as bridge formation, have been proposed (19)). Thioesters offer an experimentally tractable, prebiotically plausible reagent to model direct aminoacylation (59). By balancing substrate stability, reactivity, transfer rate and product hydrolysis, here thioesters spontaneously generate terminal aminoacylation of a majority of template-bound RNAs under near-neutral conditions in water, predominantly at the 5′ of RNAs.

Via dual-aminoacylation of junctions, this system inevitably colocalises amino acids. This could represent a candidate prebiotic opportunity to facilitate peptide production. Although aminoacyl esters exhibit a low tendency to react with amines (34), the colocalisation and inherent linkage to RNAs therein may provide fertile ground for development of ribozyme catalysts of peptide bond formation. Indeed, coupling this catalysed step to (spontaneous) direct aminoacylation and (spontaneous) peptidyl product transfer may result in iterative peptide synthesis in a nick (Fig. 5).

In this context, it is important to ask whether the 5′-ester – resistant to attack by the 2′/3′-OH and water – is also more resistant to attack by an amine of an amino acid in peptide bond formation: is the more stable 5′ form also more inert? This will depend upon the limiting step in tetrahedral intermediate formation and breakdown. The ribosome deprotonates the amine during peptide bond formation, driving productive intermediate breakdown over reversion to starting materials. If such a mechanism could be recapitulated, the nature of the hydroxyl leaving group may not be overly impactful, and prebiotic systems could effectively benefit from the stability of the 5′-aminoacyl/peptidyl ester bond.

Specific 3′-terminal overhangs drive transfers in other catalytic contexts, and we observed some aminoacyl transfer from a 3′-terminal GCCA overhang sequence, though not a 3′-terminal ACCA overhang sequence (Fig. 3*D*). Nicked loops of this nature are found in tRNAs and their minihelix analogues used in studies of synthetic RNA aminoacylation, and transfer could complicate analyses of aminoacylation of such species. As transfer would deactivate transfer RNA, the transfer capability of such overhang sequences in the acceptor stems of tRNAs merits further investigation, as it may hold implications for life’s choice of 5′-phosphorylation during tRNA biogenesis: this is established through the RNase P hydrolysis mechanism which (unusually amongst nucleolytic ribozymes) leaves a 5′-phosphate rather than the more catalytically accessible 5′-OH.

The predisposed nature of 5′-aminoacylation outlined here bridges direct chemical aminoacylation and RNA catalysis, offering opportunities for development of RNA-catalysed peptide synthesis, as well as for side-chain selectivity to connect adaptor RNA sequence to amino acid identity. Coupled to the efficiency of the flexizyme system and/or generality of thioester aminoacylation, it may also access 5′-acylated oligonucleotides to expand chemical modifications for functional RNA or potential RNA therapeutics (72, 73). Understanding the chemical and catalytic landscape of aminoacylated RNA will be critical to resolve questions of the origin and nature of biology’s translation system. The inherent biosynthetic link between 5′-and to 2′/3′-aminoacylation provides a potential bridge between primordial and modern peptide synthesis strategies.

## Materials and Methods

Standard stability assays were carried out with 0.25 µM aminoacylated FITCPriCCA oligonucleotide, incubated at 20°C or -7°C (supercooled or frozen) in 50 mM MOPS (pH 7.0 or 8.0) (@ 20°C), 200 mM NaCl, 0.05% Tween-20. Standard transfer reactions were carried out with 0.25 µM acceptor and aminoacylated donor oligonucleotides and 0.275 µM template oligonucleotide incubated at 20°C or -7°C (supercooled or frozen) in 50 mM of the stated buffer at the stated pH, 200 mM NaCl, 0.05% Tween-20. In situ aminoacylation/transfer reactions were performed at 20°C at pH 8 with 0.25 µM acceptor and unaminoacylated donor oligonucleotides and 0.275 µM template in 200 mM MOPS, 200 mM NaCl, 0.05% Tween 20. Reactions were analysed by 20% denaturing acid-PAGE, and the % hydrolysis or product was calculated by gel densitometry using ImageQuant TL software (normalised with respect to initial % aminoacylation where appropriate). All rates were calculated as initial rates, determined by fitting data to the appropriate rate equation by linear regression. Kinetic models were described by the appropriate system of differential equations and the time-dependent concentrations of all species involved were calculated using the numerical differential equation solver NDSolve in the Mathematica Software package version 14.3, using experimentally determined rate constants. Please see Supporting Text for a detailed description of compounds and methods.

## Acknowledgements and Funding Sources

This work was supported by a Royal Society University Research Fellowship (URF\R1\201271; J.A., J.M.). We would like to thank Dr Jyoti Singh for kindly supplying the thioester used in this work and Dr Benji Thoma for advice on oligonucleotide aminoacylation. We are grateful to Prof. Matthew Powner and Dr Daniel Whitaker for thoughtful suggestions on the manuscript.

