## Supplementary Information for "5′ RNA Aminoacylation via Interstrand Acyl-Transfer"

for

*James Attwater

This PDF file includes:

Supporting Text

Supporting Discussion

Supporting Note

Figures S1 to S14

Table S1

SI References

### Supporting Text

#### General Information

Reagents and solvents were purchased from Sigma-Aldrich and Fisher Scientific, unless otherwise specified. Solution pH values were measured using a Thermo Scientific Orion Star A211 Benchtop pH Meter with a Thermo Scientific Orion ROSS PerpHecT Micro Glass Bodied Combination pH Electrode. Eutectic pH values were measured using a Mettler Toledo SevenExcellence S470 pH/Conductivity Benchtop Meter with a Mettler Toledo InLab Cool Pro-ISM pH Electrode.

#### Oligonucleotide Preparation

All RNAs were polyacrylamide gel purified and precipitated in 75% EtOH (except triplets (85% EtOH) and fluorescently tagged oligonucleotides (92% EtOH)). All PCR-generated dsDNAs were purified by QIAquick PCR Purification Kit (Qiagen) prior to transcription. Oligonucleotides were quantified by measuring UV absorbance at 260 nm using a NanoDrop spectrophotometer and sequence specific extinction coefficients derived from OligoCalc (1).

#### PAGE Analysis

All gels were scanned using the appropriate laser wavelength on a Typhoon FLA-9000 imager (Cytiva) and quantified by gel densitometry using ImageQuant TL (Cytiva) and defining a manual background between the minima in between bands. Species % was obtained by same-lane normalised band intensity.

#### Flexizyme Aminoacylation Reactions:

For preparative aminoacyl RNA synthesis, the appropriate flexizyme and the substrate oligonucleotide (each 10 μM, 0.6-1 nmol) were annealed (94°C for 2 min) in HEPES (final concentration 50 mM, pH 7.5 or 8) before being made up to 600 mM $MgCl_2$ (from 3 M stock) and 5 mM amino acid substrate (dissolved in DMSO-$d_6$; from 25 mM stock). Reactions were incubated on ice for between 2 and 5 h and stopped with the addition of chilled NaOAc (pH 5.2) to a final concentration of 300 mM. Reactions were then precipitated on ice in 75% or 92% EtOH, before being resuspended in 5 µL NaOAc (pH 5.2) and added to 30 µL acid loading buffer (ALB; 93% formamide, 150 mM NaOAc, 10 mM EDTA; pH 5.2). Products were separated by 6 M urea EDTA-Acetate buffer (EA; 100 mM NaOAc, 10 mM EDTA; pH 5.2) denaturing acid-PAGE (20% 19:1 acrylamide:bis-acrylamide; run chilled at 4°C) and excised using fluorescence or UV transillumination. Products were then eluted from gel fragments in 0.3 M NaOAc (pH 5.2) on ice, filtered with Spin-X column filters and precipitated in 75% or 92% ethanol before being resuspended in water or 1 mM NaOAc (pH 5.2) and quantified. Aminoacylated RNAs were stored at -70°C.

Analytical flexizyme reactions were carried out according to the same protocol but at smaller scale (oligonucleotides each 1 µM, 5 pmol). Timepoints were taken (0.5 µL, 0.5 pmol), added to 6 µL ALB and analysed by denaturing acid-PAGE. Gels were scanned and bands quantified by densitometry.

#### Stability Incubations:

Incubations were carried out in the conditions stated for the specific reaction. Generally, aminoacylated RNA (0.25 µM) was incubated in buffer at a set temperature and 0.5 pmol timepoints were taken at regular intervals. These were added to a 6-fold volume of ALB (and a 30-fold excess of the appropriate template-binding competitor RNA if a template strand was present) and stored at -70°C until being analysed by denaturing acid-PAGE. Gels were scanned and bands quantified by densitometry.

#### Transfer Reactions:

Reactions were carried out in the conditions stated for the specific reaction. Generally, acceptor and template oligonucleotides (0.25 µM final concentration, 10% excess template) were mixed in buffer before adding the aminoacylated donor oligonucleotide (to 0.25 μM). The reactions were then incubated at a set temperature (after first flash freezing in liquid nitrogen for frozen reactions). Timepoints were taken, added to a six-fold excess of ALB, and a 30-fold excess of template-binding competitor RNA, and analysed by denaturing acid-PAGE. Gels were scanned and bands quantified by densitometry.

#### Thioester Transfer Reactions:

Reactions with in situ thioester aminoacylation were carried out in the conditions stated for the specific reaction. Generally, all oligonucleotides (0.25 μM, 10% excess template) were mixed in buffer before L-Ala-SEt·HCl* (200 mM, pH adjusted to 7.0 with NaOH/HCl) was added. Reactions were then incubated at constant and timepoints were taken at regular intervals. These were added to a six-fold excess of ALB with a 30-fold excess of template-binding competitor RNA and analysed by denaturing acid-PAGE. Gels were scanned and bands quantified by densitometry.

#### 5'-Aminoacyl RNA Synthesis:

5'-aminoacyl RNA was synthesised using scaled up thioester transfer reactions (oligonucleotides at 2 μM, 400 pmol, 10% excess template). Reactions were flash frozen in liquid nitrogen then incubated at -7°C for 48 h. Reactions were precipitated in 300 mM NaOAc (pH 5.2), 92% EtOH and resuspended in template-binding competitor RNA (200 μM, 30-fold excess) and 60 μL ALB and products were separated by denaturing acid-PAGE. Bands were excised using UV transillumination and products eluted from gel fragments in 0.3 M NaOAc (pH 5.2) on ice, filtered with Spin-X column filters and precipitated in 92% ethanol before being resuspended in water or 1 mM NaOAc (pH 5.2) and quantified. Aminoacylated RNAs were stored at -70°C.

---

*The aminoacyl thioester used in these experiments (L-Ala-SEt) was kindly donated by Dr Jyoti Singh in the Powner group. Synthesis and characterisation are given in ref. (13).

#### Stability Kinetics:

The observed fraction of 2'/3'-aminoacylation at each timepoint ($P_t^3$) was normalised ($P_N^3$) with respect to that at time $t = 0$ ($P_0^3$), $P_N^3 = \frac{P_t^3}{P_0^3}$. The fraction of 2'/3'-aminoacylation was then plotted as $-\ln(P_N^3)$ against $t$ and the observed rate constant ($k_{hyd}^3$) determined by fitting the data to the pseudo-first-order exponential decay by linear regression, according to the equation $P_N^3 = e^{-k_{hyd}^3 t}$. From this the half-life ($t_{1/2}$) was calculated by $t_{1/2} = \frac{\ln(2)}{k_{hyd}^3}$. The hydrolysis rate for 5'-aminoacyl species ($k_{hyd}^5$) was determined analogously from purified starting material.

#### Transfer Kinetics:

The observed fraction of 5'-aminoacylation ($P_t^5$)) at each timepoint was normalised ($P_N^5$) with respect to the initial fraction of 3'-aminoacylation on the donor oligonucleotide, $P_N^5 = \frac{P_t^5}{P_0^3}$. The fraction of 5'-aminoacylation was then plotted as $-\ln(1 - P_N^5)$ against $t$ andthe observed rate constant ($k_{tr}$) determined by fitting the early data to the first order rate equation $1 - P_N^5 = e^{-k_{tr}t}$ by linear regression. As both $k_{hyd}^3$ and $k_{hyd}^5 \ll k_{tr}$, at early timepoints $1 - P_N^5 \approx P_N^3$.

#### Thioester Kinetics:

$P_t^3$ at each timepoint was determined by subtracting the total level of internal aminoacylation in the negative control from the total level of aminoacylation in the reaction (Fig. S12). The fraction of aminoacylation was then plotted as $-\ln(1 - P_t^3)$ against $t$. The observed pseudo-first order rate constant ($k_{obs}$) was then determined by fitting the early data to the first order rate equation $1 - P_t^3 = e^{-k_{obs}t}$ by linear regression. The reaction can be treated as first order as $[thioester] \gg [oligonucleotides]$, however, due to the hydrolysis of the product, $k_{obs}$ is measured as the initial rate from early timepoints.

#### Kinetic Models:

Full reaction systems (Fig. 4c, lower panel) were modelled according to the following differential equations:

$$\frac{dP^3}{dt} = -k_{tr}P_t^3 - k_{hyd}^3 P_t^3 + k_{hyd}^5 P_t^D + k_{obs}P_t^U + k_{rev}P_t^5$$

$$\frac{dP^5}{dt} = k_{tr}P_t^3 - k_{hyd}^5 P_t^5 - k_{obs}P_t^5 + k_{hyd}^3 P_t^D - k_{rev}P_t^5$$

$$\frac{dP^U}{dt} = k_{hyd}^3 P_t^3 + k_{hyd}^5 P_t^5 - k_{obs}P_t^U$$

$$\frac{dP^D}{dt} = k_{obs}P_t^5 - k_{hyd}^3 P_t^D - k_{hyd}^5 P_t^D$$

Where:

$P^3$ = the fraction of nicks that are aminoacylated at the 3'-terminus

$P^5$ = the fraction of nicks that are aminoacylated at the 5'-terminus

$P^U$ = the fraction of nicks that are unaminoacylated

$P^D$ = the fraction of nicks that are aminoacylated at both the 5'- and 3'-terminus (dual aminoacylated)

$k_{hyd}^3$ = the rate of hydrolysis of 3'-aminoacylation

$k_{hyd}^5$ = the rate of hydrolysis of 5'-aminoacylation

$k_{tr}$ = the rate of forward (3' → 5') transfer

$k_{rev}$ = the rate of reverse (5' → 3') transfer

$k_{obs}$ = the rate of 3'-aminoacylation

Simplified model systems (Fig. 4a, Fig. 4c, upper panel) were modelled using the same equations less any terms containing rates or fractions omitted from that specific system.

#### Dephosphorylation of Triphosphorylated Oligonucleotides:

Shrimp Alkaline Phosphatase (rSAP; NEB; 5 units) was added to a reaction of 1 × rCutSmart Buffer (NEB) and pppCCA (5 nmol, 50 μM) on ice. The reaction was mixed by pipette and incubated at 37°C for 8 h, then 65°C for 5 minutes. To the reaction was added urea (9 M, 206 μL) and EDTA (0.5 M, 3 μL) and purified by denaturing PAGE (30% 19:1 acrylamide:bis-acrylamide), next to a reference lane of starting material. The product band was excised using UV transillumination, eluted from gel fragments in NaOAc (0.3 M, pH 5.2), filtered and precipitated in 85% ethanol before being resuspended in water and quantified.

#### Phosphorylation of cy5PriG:

T4 Polynucleotide Kinase (PNK; NEB; 30 units) was added to a reaction of 1 × PNK Buffer (NEB), ATP (1 mM) and Cy5PriG (1 nmol, 6.67 μM) on ice. The reaction was mixed by pipette and incubated at 37°C for 0.5 h. To the reaction was added NaOAc (15 μL, 3 M, pH 5.2) and the mixture was precipitated in and washed with 92% EtOH. The pellet was resuspended in H2O (20 μL), urea (9 M, 40 μL) and EDTA (0.5 M, 3 μL) and purified by denaturing PAGE, next to a reference lane of starting material. The product band was excised using UV transillumination, eluted from gel fragments in NaOAc (0.3 M, pH 5.2), filtered and precipitated in 92% ethanol before being resuspended in water and quantified.

#### Synthesis of L-Alanine-DBE

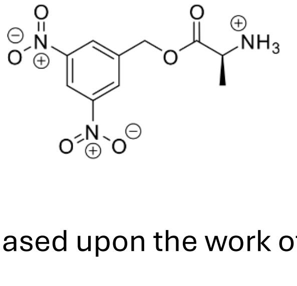

L-Alanine-DBE was synthesised based upon the work of Suga et al. (2). To a solution of *N*-Boc-alanine (260 mg, 1.2 mmol) in DMF (200 μL) and $NEt_3$ (278.2 μL, 2 mmol) was added 3,5-dinitrobenzyl chloride (216.58 mg, 1 mmol) and stirred at room temperature overnight. Reaction was diluted with $Et_2O$ (9 mL). The liquid phase was washed with HCl (0.5 M, 6 mL × 3), saturated $NaHCO_3$ (6 mL × 3) and brine (10 mL), dried over $MgSO_4$ and concentrated under vacuum to achieve crude product as a gummy yellow solid (302 mg, 0.82 mmol, 68% yield). To a stirred solution of *N*-Boc-L-alanine-DBE (302 mg, 0.82 mmol) in $Et_2O$ (1 mL) was added HCl (2M in $Et_2O$; 12 mL) slowly at 0°C. The mixture was stirred for 2 days at room temperature. The reaction mixture was concentrated under reduced pressure to provide a gummy liquid. This was washed with $Et_2O$ (5 mL × 3) to yield the product hydrochloride salt as a white powder (245 mg, 0.80 mmol, 67% overall yield). $^1H$ NMR (600 MHz, DMSO-$d_6$): $\delta_H$ 8.83 (1H, t, $CHC(NO_2)C\mathit{H}C(NO_2)$), 8.75 (2H, d, $CH_2CC\mathit{H}C(NO_2)$), 8.54 (3H, br s, $\mathit{H_3}NCH(CH_3)C(O)$), 5.50 (2H, s, $OC\mathit{H_2}C$), 4.25 (1H, q, $H_3NC\mathit{H}(CH_3)C(O)$), 1.46 (3H, d, $H_3NCH(C\mathit{H_3})C(O)$).

#### Synthesis of D-Alanine-DBE

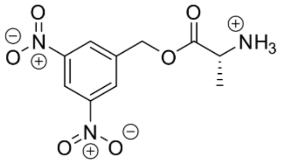

D-Alanine-DBE was synthesised based upon the work of Suga et al. (2). To a solution of *N*-Boc-D-alanine (260 mg, 1.2 mmol) in DMF (200 µL) and $NEt_3$ (278.2 µL, 2 mmol) was added 3,5-dinitrobenzyl chloride (216 mg, 1 mmol) and stirred at room temperature overnight. Reaction was diluted with $Et_2O$ (9 mL). The liquid phase was washed with HCl (0.5 M, 6 mL × 3), saturated $NaHCO_3$ (6 mL × 3) and brine (10 mL), dried over $MgSO_4$ and concentrated under vacuum to achieve crude product as gummy yellow solid (336 mg, 0.91 mmol, 76% yield). *N*-Boc-D-alanine-DBE (336 mg, 0.91 mmol) was dissolved in HCl (2M in $Et_2O$; 4 mL), added dropwise at 0°C. The mixture was stirred for 4 days at room temperature, with a further 2 mL of HCl (2M in $Et_2O$) added after 2 days. The reaction mixture was concentrated under reduced pressure to provide a gummy liquid. This was washed with $Et_2O$ (10 mL × 3) to yield the product as a hydrochloride salt as a white powder (265.6 mg, 0.869 mmol, 72% overall yield). $^1H$ NMR (500 MHz, DMSO-$d_6$):

$\delta_H$ 8.82 (1H, t, CHC(NO$_2$)C*H*C(NO$_2$)), 8.75 (2H, d, CH$_2$CC*H*C(NO$_2$)), 8.66 (3H, br s, *H$_3$*NCH(CH$_3$)C(O)), 5.49 (2H, s, OC*H$_2$*C), 4.22 (1H, q, H$_3$NC*H*(CH$_3$)C(O)), 1.47 (3H, d, H$_3$NCH(C*H$_3$*)C(O)).

#### Synthesis of L-Phenylalanine-CME

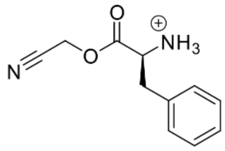

L-Phenylalanine-CME was synthesised based upon the work of Karmakar et al. (3). To a solution of tert-butoxycarbonyl-phenylalanine (1 g, 3.77 mmol) in DMF (20 mL) and NEt$_3$ (1.05 mL, 7.54 mmol) at 0°C was added chloroacetonitrile (240 μL, 3.77 mmol). It was stirred at room temperature for 2h. Reaction was quenched with ice and compound was extracted with EtOAc (4 × 25 mL). Combined organic layer was washed with brine (50 mL), dried over Na$_2$SO$_4$ and concentrated under vacuum (co-evaporated with heptane) to achieve crude product. It was later purified by column chromatography (SiO$_2$; EtOAc/pet. Ether, 1:3) to achieve white solids of *N*-Boc-L-phenylalanine-CME (0.166 g, 0.546 mmol, 15% yield). To a stirred solution of *N*-Boc-phenylalanine-CME (0.166 g, 0.546 mmol) in MeCN (4 mL) was added HCl (4 M in dioxane) (273 μL, 1.092 mmol) slowly at 0°C. The mixture was stirred for 2 h. During this period reaction temperature was maintained at 10-15°C. The reaction mixture was concentrated at 30-35°C under reduced pressure to provide a gummy liquid. The gummy crude was dissolved in minimum amount of MeCN and to that solution was added excess Et$_2$O, precipitating out the desired product as white solids. The solids were filtered, washed with Et$_2$O & dried under vacuum to produce desired product as hydrochloride salt as a white powder (0.0722 g, 0.354 mmol, 65% yield). $^1$H NMR (500 MHz, DMSO-d$_6$): $\delta_H$ 8.60 (3H, br s, N*H$_3$*CH(CH$_2$C$_6$H$_5$)C(O)), 7.30 (5H, m, NH$_3$CH(CH$_2$C$_6$*H$_5$*)C(O)), 5.09 (1H, d, J=16.1 Hz, OC*H*HCN), 5.07 (1H, d, J=16.1 Hz, OCH*H*CN), 4.40 (1H, t, NH$_3$C*H*(CH$_2$C$_6$H$_5$)C(O)), 3.20 (1H, dd, J=14.1, 5.9 Hz, NHCH(C*H*HC$_6$H$_5$)C(O)), 3.10 (1H, dd, J=14.1, 7.6 Hz, NHCH(CH*H*C$_6$H$_5$)C(O)).

#### Synthesis of *N*-Biotinyl-L-Phenylalanine-CME

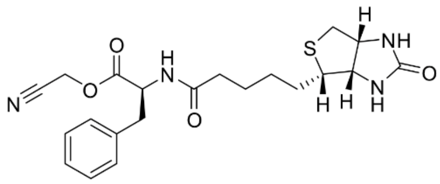


To synthesise *N*-Biotinyl-L-Phenylalanine Cyanomethyl Ester, L-Phenylalanine (247.8 mg, 1.5 mmol) was dissolved in NaOH (1 M, 1.375 mL) and cooled to 4°C, it was then mixed with NHS-Biotin (512.1 mg, 1.5 mmol) dissolved in DMSO (2.143 mL) and the mixture incubated at 4°C overnight. HCl (2 M, 1.440 mL) was added to the reaction mixture, the resulting gel was transferred to a filter paper and washed with HCl (0.1 M, 5 mL × 3) and water (3 mL), then dried *in vacuo* overnight. The product was obtained as a white solid, dissolved in MeOH (200 mL) with triethylamine (90 μL, 65.34 mg, 0.65 mmol) and purification was attempted by column chromatography (SiO2; EtOAc (1% AcOH)/MeOH, 8:2). This yielded 620 mg of a mixture of product, DMSO and trace quantities of starting material as a white powder. Crude *N*-biotinyl-L-phenylalanine (587.25 mg, 1.5 mmol (assuming 100% yield up to this point)) was mixed with triethylamine (2.5 mL, 17.9 mmol) and dissolved in DMF (3 mL). Chloroacetonitrile (189.8 μL, 3mmol) was added, and the reaction mixture stirred at room temperature overnight. The reaction mixture was concentrated under reduced pressure and the DMF co-evaporated with toluene. The crude product was then redissolved in DMF (25 mL) and diluted with EtOAc (250 mL), then washed with water (100 mL × 2) and brine (100 mL). The organic layer was dried over $MgSO_4$ and concentrated under reduced pressure. The aqueous layers were extracted with EtOAc (100 mL × 2), the organic layers were combined and washed with brine (100 mL) then dried over $MgSO_4$ and concentrated under reduced pressure. The product was obtained as a light-yellow powder (355.3 mg, 0.825 mmol, 55% yield). $^1H$ NMR (500 MHz, DMSO-$d_6$): $\delta_H$ 8.39 (1H, d, CHN*H*C(O)$CH_2$), 7.25 (5H, m, $CH_2C_6H_5$), 6.38 (1H, s, CHN*H*C(O)NHCH), 6.34 (1H, s, CHNHC(O)N*H*CH), 4.98 (1H, d, J=16.0 Hz, OC*H*HCN), 4.97 (1H, d, J=16.0 Hz, OCH*H*CN), 4.52 (1H, m, NHC*H*($CH_2C_6H_5$)C(O)), 4.30 (1H, dd, J=7.7, 5.3 Hz, NHC*H*$CH_2$S), 4.10 (1H, m,

NHC*H*CH($CH_2$)S), 3.10 (2H, m, C*H*H($C_6H_5$) and NHCHC*H*($CH_2$)S overlapping), 2.92 (1H, dd, J=13.7, 9.6 Hz, CH*H*($C_6H_5$)), 2.82 (1H, dd, J=12.5, 5.1, SC*H*HCH), 2.58 (1H, d, J=12.5 Hz, SCH*H*CH), 2.06 (2H, m, $CH_2$C*H₂*C(O)NH), 1.56 (1H, m, NHC(O)$CH_2CH_2CH_2$C*H*HCH), 1.42 (3H, m, NHC(O)$CH_2CH_2CH_2$CH*H*CH and NHC(O)$CH_2$C$H_2$$CH_2CH_2$CH overlapping), 1.22 (2H, m, NHC(O)$CH_2CH_2$C$H_2$$CH_2$CH).

#### Supporting Discussion

The oligonucleotide-aminoacyl ester hydrolysis rate in ice behaves consistent with a linear relationship to [$^{-}$OH], after accounting for predicted cooling-induced increases in MOPS buffer pH (from 7 to 7.16 and from 8 to 8.46). In pH 7 buffers, the half-life of alanyl RNA ester increased to over 8 days when in supercooled solution at -7°C vs 20 h at 20°C. Freezing such supercooled solutions then halved stability at -7ºC, likely due to the waning influence of polyelectrolyte effects in the salty eutectic phase (4, 5) in which solutes were concentrated 5-fold (see Supporting Note) (6, 7). We also tested the influence of $Mg^{2+}$ cations (necessary for ribozyme structure and catalysis (8, 9) and relied on by the most ancient parts of the ribosome (10)) whose reported effects upon aminoacyl ester stability vary across the literature (4, 11, 12). We observed a modest 1.6-fold destabilisation of the aminoacyl bond by $Mg^{2+}$ (pH 7, -7°C frozen), while a larger 2.6-fold destabilisation was observed in supercooled solutions (Fig. S2), presumably due to a larger relative increase in ionic strength.

#### Supporting Note

The increase in concentration of the eutectic phase was determined according to previously reported methods (6). Eutectic phase volume ($V_e$) at -7°C was estimated by assessing persistence of solid ice at various reaction concentrations. 'Dummy' 50 µl transfer reactions, with standard concentrations of all components other than oligonucleotides were first lyophilized. The resulting salts were then resuspended in a range of volumes of water below 50 µl ($V_n$). These higher concentration samples were refrozen at -70°C, before being transferred to -7°C to equilibrate for 48 h. The melting of solid ice upon transfer to -7°C was taken to show the new solute concentration ($[S]_n$) was higher than the eutectic solute concentration ($[S]_e$), lowering the solution's freezing point to below that of the original eutectic phase, and hence that $V_n < V_e$. Conversely, the persistence of solid ice upon transfer to -7°C showed that $[S]_n < [S]_e$, elevating the freezing point of the solution and implying $V_n > V_e$. The inflection point between ice persistence and ice melting was observed around $V_n$ = 10 µl. $V_e$ is therefore, ~20% of initial reaction volume. This represents a 5-fold concentration of solutes in the eutectic phase and reveals the eutectic phase to be around 1 M $MgCl_2$, 250 mM MOPS, 0.25 % Tween-20, and 1.25 µM of each RNA.

### Supporting Figures

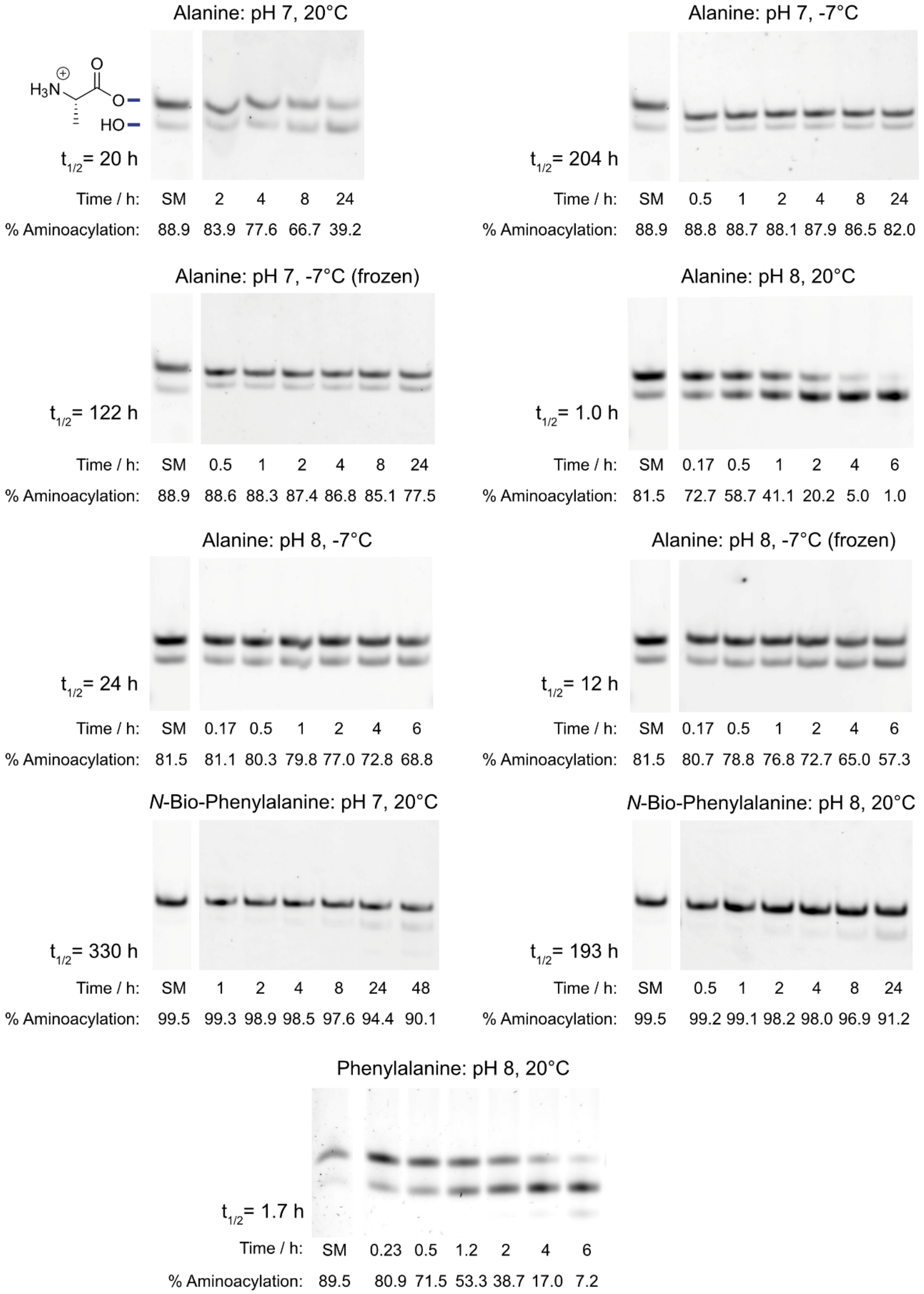


**Fig. S1 | The effect of pH and temperature on the stability of aaRNA.** PAGE scans of stability assays for aminoacyl esters of alanine and *N*-Bio-phenylalanine and phenylalanine at pH 7 and 8 at different temperatures. Aminoacyl RNA (0.25 µM) was incubated in MOPS (50 mM, pH 7.0 or 8.0 (@ 20°C)), NaCl (200 mM), Tween-20 (0.05%) at 20°C or -7°C (supercooled or frozen). Amino acid, pH and temperature stated above, and half-life stated to the left of each scan, ‘SM’ denotes starting material.

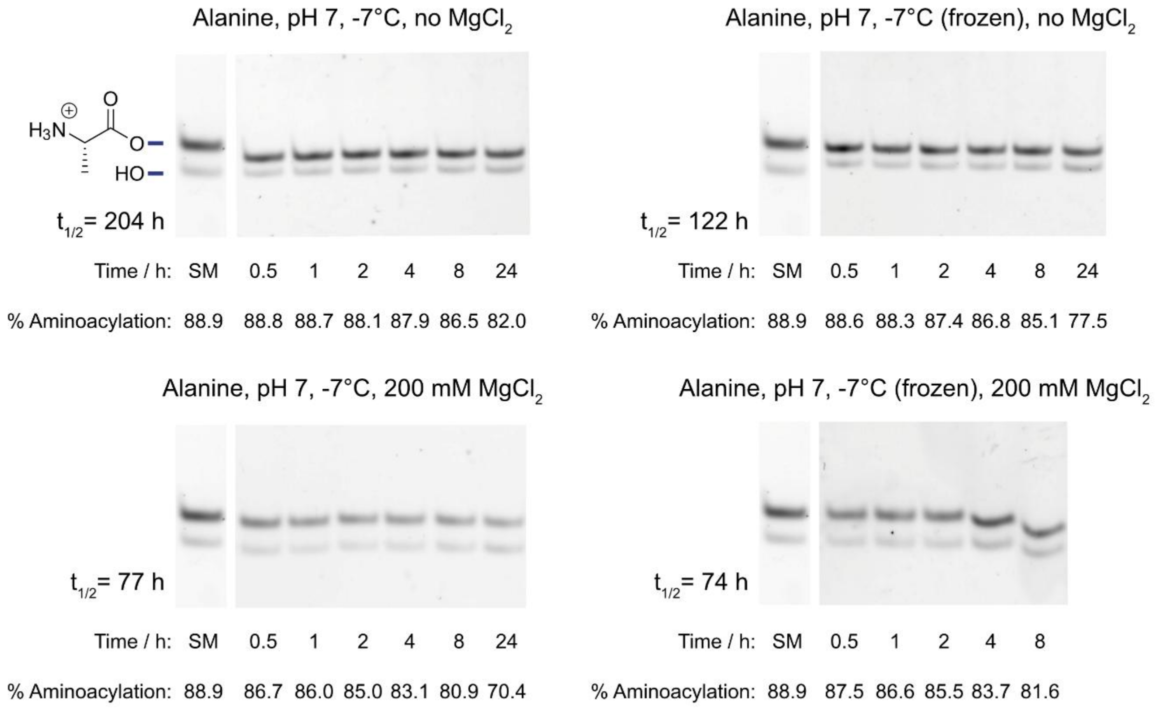


**Fig. S2 | The effect of the presence of magnesium ions on aminoacyl stability at -7°C in supercooled or frozen conditions**. Above: PAGE scans of alanine stability assays with no $MgCl_2$ present (from Fig.S1). Below: PAGE scans of the same conditions but with buffers also containing 200 mM $MgCl_2$. The destabilising effect in supercooled conditions (2.6-fold) is higher than that in frozen conditions (1.6-fold). Half-lives are given to the left of each scan, 'SM' denotes starting material.

a

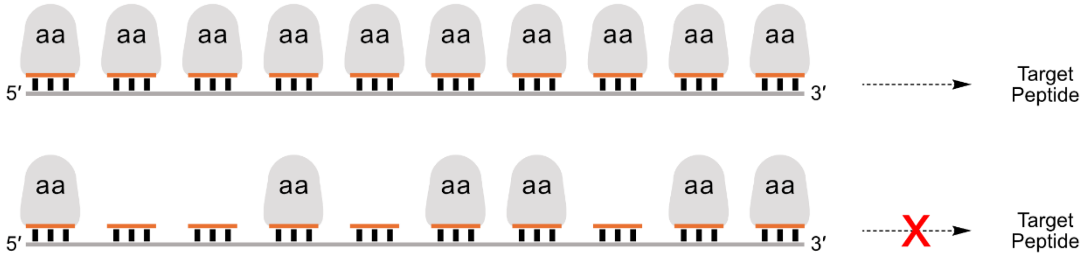


b

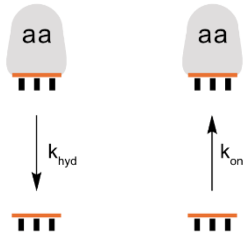


c

| $k_{hyd}$ ($h^{-1}$) | $k_{on}$ ($h^{-1}$) | Steady State % aa | % Complete 10-mer |
|---|---|---|---|
| 0.73 | 0.15 | 17 | $2.0 \times 10^{-6}$ |
| 0.73 | 0.73 | 50 | 0.1 |
| 0.73 | 1.27 | 64 | 1.2 |
| 0.058 | (0.15) | 72 | 3.7 |
| 0.058 | (0.73) | 93 | 48 |
| 0.058 | (1.27) | 96 | 64 |

**Fig. S3 | Toy model exploring how different steady state aminoacylation levels translate into successful 10-mer peptide synthesis. a**, Schematic of adapter RNAs on a template ‘proto mRNA’ encoding a 10-mer peptide product. All adapter RNAs on the proto mRNA are required to be aminoacylated in order to synthesise the desired peptide. **b,** Adapter RNAs are in flux, with hydrolysis at rate $k_{hyd}$ removing aminoacylation and chemical or ribozymatic aminoacylation reinstating it at rate $k_{on}$. **c**, Table of the outcomes of various $k_{hyd}$ and $k_{on}$ rates. The two hydrolysis rates chosen are those observed for 2'/3'-Ala at pH 8, 20°C (0.73 $h^{-1}$) and at pH 8, -7°C (frozen) (0.058 $h^{-1}$). Aminoacylation rates were chosen to illustrate different levels of steady state aminoacylation (‘Steady State % aa’ in table) and the corresponding levels of template fully covered in translation-competent adapters. Aminoacylation rates for rows where $k_{hyd}$= 0.058 $h^{-1}$ are shown in parentheses to indicate the likelihood of correspondingly lower $k_{on}$ rates at lower temperatures.

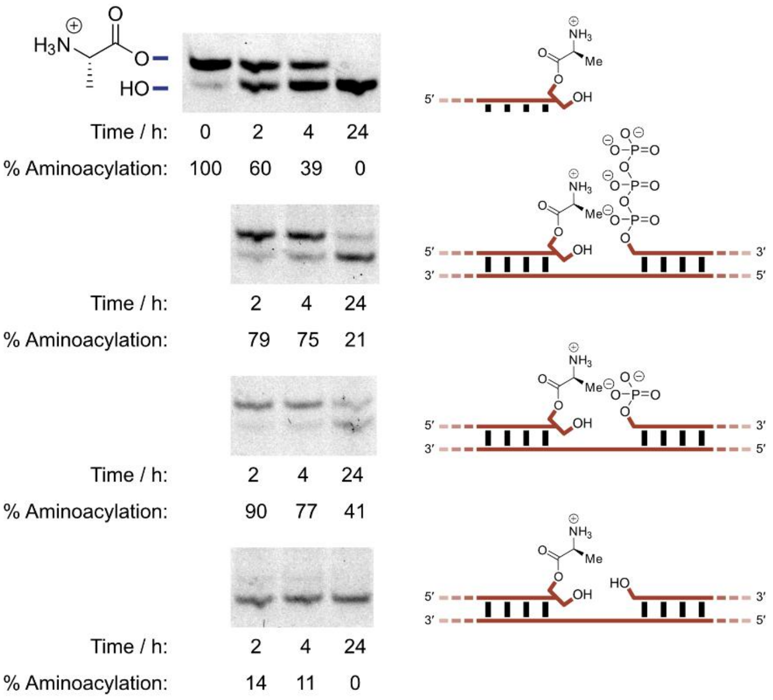


**Fig. S4 | The effect of adjacent phosphate groups on the stability of aaRNA.**
FITCPriCCA-Ala (0.25 µM) was incubated alone or with template (TempCCACCC, 0.3 µM) and triplets (pppCCC, pCCA or HO-CCA, 2 µM) in HEPBS (50 mM, pH 8.3), $MgCl_2$ (200 mM) at -7°C (frozen).

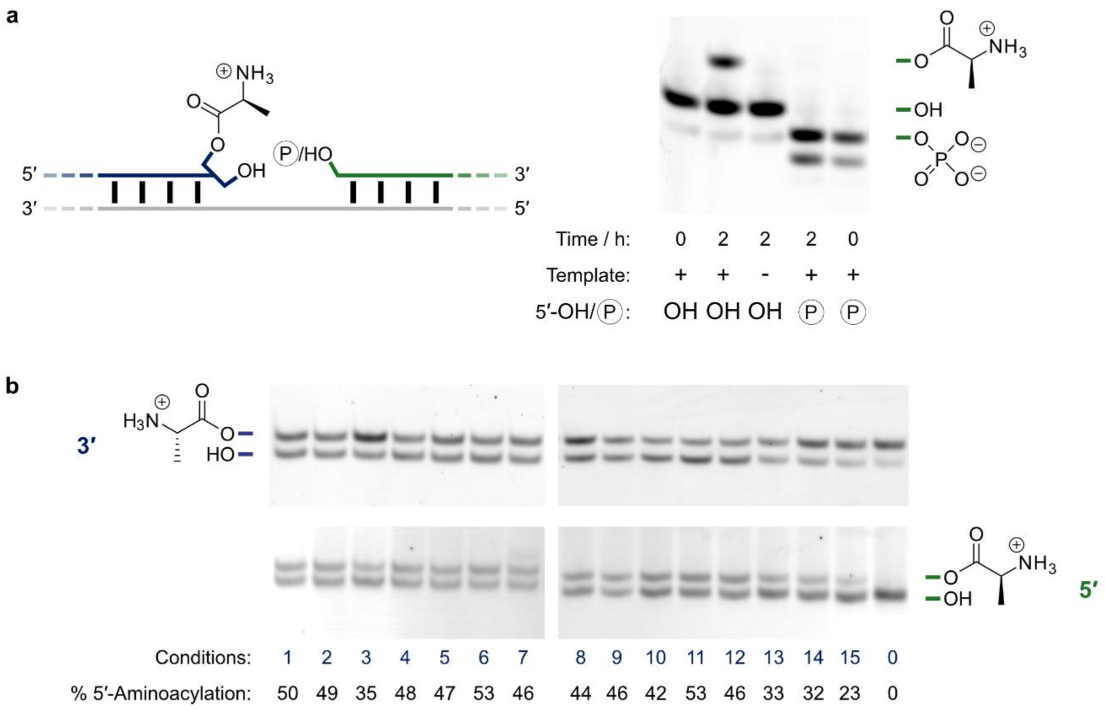


Conditions:

| Buffer | pH | Conc. (mM) | Salts: $MgCl_2$ 200mM | $MgCl_2$ 10 mM | NaCl 200 mM | KCl 200 mM | $MgCl_2$ & KCl 100 mM | $MgCl_2$ & NaCl 100 mM | None |
|---|---|---|---|---|---|---|---|---|---|
| HEPBS | 8.3 | 50 | 1 | 3 | 6 | 7 | 5 | 4 | 2 |
| | | 100 | 8 | | | | | | |
| | 8.6 | 50 | 9 | | | | | | |
| | 9.0 | 50 | 10 | | | | | | |
| TRIS | 8.3 | 50 | 11 | | | | | | |
| | 7.9 | 50 | 12 | | | | | | |
| | 7.4 | 50 | 13 | | | | | | |
| HEPES | 7.5 | 50 | 14 | | | | | | |
| MES | 7.0 | 50 | 15 | | | | | | |

**Fig. S5 | The transfer reaction is dependent on the presence of both 5'-hydroxyl group and template strand and occurs in a broad range of buffers and salt conditions. a**, Transfer reactions (3'-Ala-FITCPriCCA (0.25 µM), Cy5PriG (0.25 µM) or pCy5PriG (0.25 µM), TempG (0.275 µM), MOPS (50 mM, 8.0 (@ 20°C)), NaCl (200 mM), Tween-20 (0.05%)) were set up with and without template and 5'-OH. Reactions without 5'-OH contain 5'-monophosphate Cy5PriG (pCy5PriG), shown as an encircled ‘P’. Reactions were incubated at 20°C for 2 h, initial timepoints are included to show the relative mobility of Cy5PriG and pCy5PriG. 5'-aminoacylation is only observed in the presence of both template and 5'-OH. **b**, Transfer reactions work in a range of salt and buffer conditions (see lower table; lane 0 = starting material; 0.25 µM FITCPriCCA-Ala, 0.275 µM sTQT2, 0.25 µM HO10Cy5, 0.05% Tween-20, incubated at -7°C (frozen) for 3h).

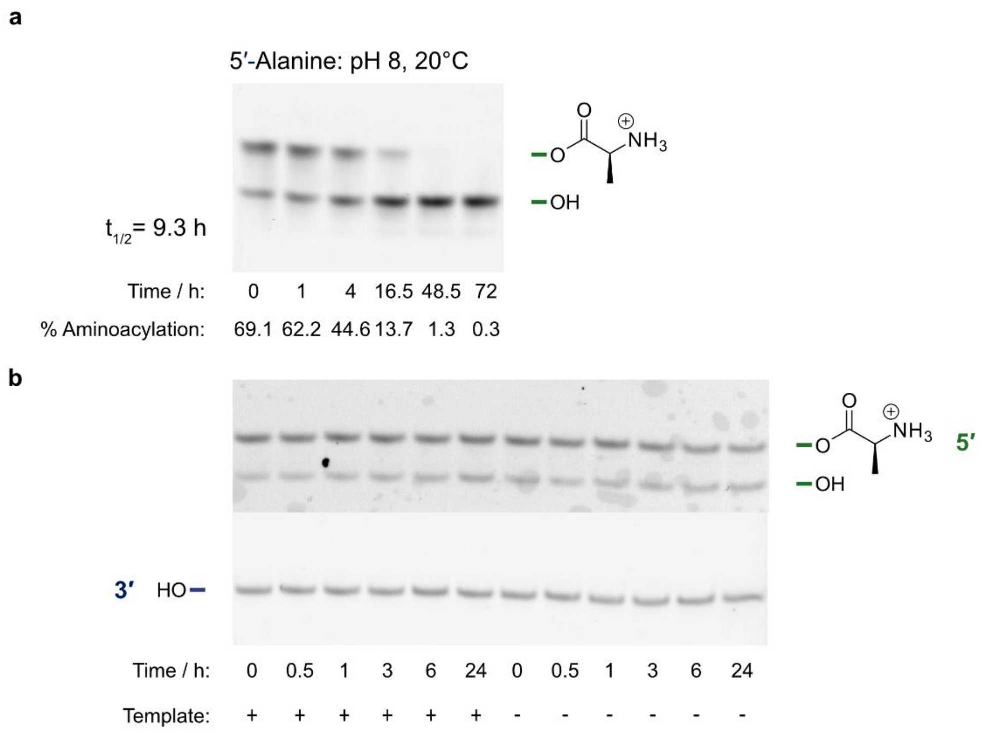


**Fig. S6 | 5'-aminoacylation is much more stable and the reverse transfer is not observed. a,** PAGE scan of stability assay for 5'-aminoacyl ester of alanine. Aminoacyl RNA (0.25 µM) was incubated in MOPS (50 mM, 8.0 (@ 20°C)), NaCl (200 mM), Tween-20 (0.05%) at 20°C. **b**, PAGE scan of 5'-Ala Cy5PriA (0.25 µM) incubated with FITCPriCCA (0.25 µM), with or without TempA (0.275 µM) in MOPS (pH 8, 50 mM), NaCl (200 mM), Tween-20 (0.05%) at -7°C (frozen). No 3'-aminoacylation band is observed; after 24 h under these conditions, the corresponding 2'/3'- to 5'-transfer would be complete, and 2'/3'-hydrolysis would be mostly complete. The average background signal when quantifying where the 3'-aminoacylation band would appear is 0.082% ± 0.036%, giving a detection limit of 0.155% (background + 2σ). Using our kinetic model (Fig. 4) a backward (5'-3') rate of 0.0098 $h^{-1}$ limits the level of 3'-aminoacylation to below this detection limit.

a

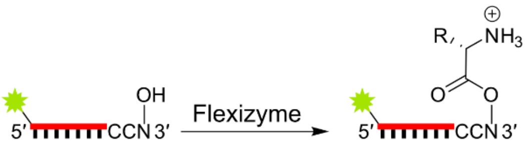


b

| Flexizyme | Amino Acid Substrate | Oligonucleotide Substrate | pH | Time /h | % Aminoacylation |
|---|---|---|---|---|---|
| eFx | L-Phe-CME | FITCPriCC**A** | 7.5 | 2 | **51** |
| eFx | L-Phe-CME | FITCPriCC**A** | 7.5 | 5 | **70** |
| dFx | L-Ala-DBE | FITCPriCC**A** | 7.5 | 2 | **27** |
| dFx | L-Ala-DBE | FITCPriCC**A** | 7.5 | 5 | **48** |
| eFx | L-Phe-CME | FITCPriCC**C** | 7.5 | 2 | **17** |
| eFx | L-Phe-CME | FITCPriCC**C** | 7.5 | 6.5 | **28** |
| dFx | L-Ala-DBE | FITCPriCC**C** | 7.5 | 2 | **1.0** |
| dFx | L-Ala-DBE | FITCPriCC**C** | 7.5 | 6.5 | **3.1** |
| eFx | L-Phe-CME | FITCPriCC**G** | 8 | 1.5 | **9.8** |
| eFx | L-Phe-CME | FITCPriCC**G** | 8 | 3.75 | **38** |
| dFx | L-Ala-DBE | FITCPriCC**G** | 8 | 1.5 | **15** |
| dFx | L-Ala-DBE | FITCPriCC**G** | 8 | 3.75 | **24** |
| eFx | L-Phe-CME | FITCPriCC**U** | 8 | 1.5 | **18** |
| eFx | L-Phe-CME | FITCPriCC**U** | 8 | 3.75 | **26** |
| dFx | L-Ala-DBE | FITCPriCC**U** | 8 | 1.5 | **1.7** |
| dFx | L-Ala-DBE | FITCPriCC**U** | 8 | 3.75 | **1.9** |
| - | L-Phe-CME | FITCPriCC**A** | 7.5 | 5 | **1.2** |
| - | L-Phe-CME | FITCPriCC**C** | 7.5 | 6.5 | **1.9** |
| - | L-Phe-CME | FITCPriCC**G** | 8 | 3.75 | **6.2** |
| - | L-Phe-CME | FITCPriCC**U** | 8 | 3.75 | **6.5** |

**Fig. S7 | The use of flexizymes to aminoacylate RNA of terminus -CCN. a**, Schematic of flexizyme reaction. **b**, Flexizyme reactions were carried out with eFx or dFx (1 μM), FITCPriCCN (1 μM), L-Phe-CME or L-Ala-DBE (5 mM) in HEPES (pH 7.5, 50 mM) or MOPS (pH 8, 50 mM), $MgCl_2$ (600 mM), DMSO (20%) at 0°C. Higher pH was found to increase yield in some cases. Background reactivity for L-Phe-CME is shown, no background reactivity was observed for L-Ala-DBE under reaction conditions.

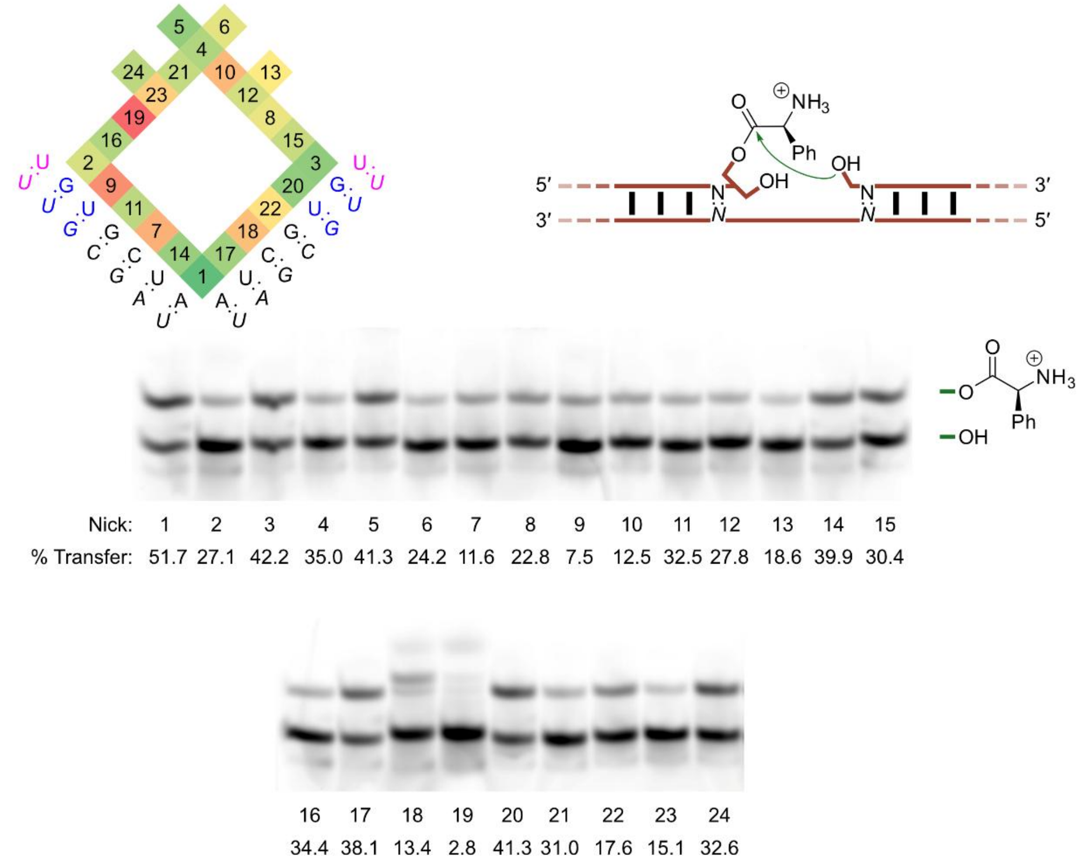


**Fig. S8 | The transfer reaction is dependent on the sequence of the nick.** 24 combinations (Fig. 3b) of template (0.275 µM of individual TempN/NN RNAs, see Oligonucleotide Sequences), donor and acceptor strands (0.25 µM of individual FITCPriCCN/Cy5PriN RNA combinations) exhibited different levels of transfer in MOPS (pH 8, 50 mM), NaCl (200 mM), Tween-20 (0.05%) at -7°C (frozen) after 24 h. % transfer is normalised to the starting level of 3'-aminoacylation.

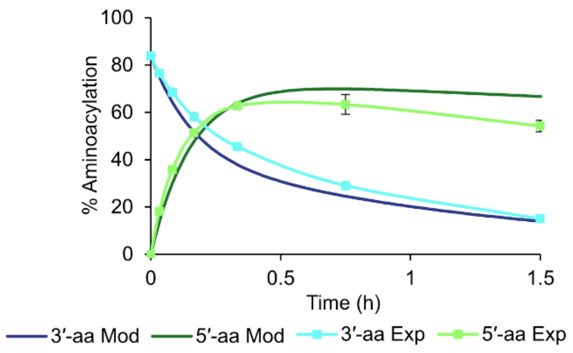


**Fig. S9 | Modelled transfer recapitulates the accumulation of 5'-aminoacylation seen in experimental data.** The model levels of 3'- and 5'-aminoacylation (darker colours) match well with those observed (lighter colours). The experimental data is for the transfer reaction carried out at pH 8 as in Fig. 1c, the model shown uses rates calculated in these conditions, the same initial level of 3'-aminoacylation is used and a two-fold excess of FITCPriCCA is modelled, as used in the experiment.

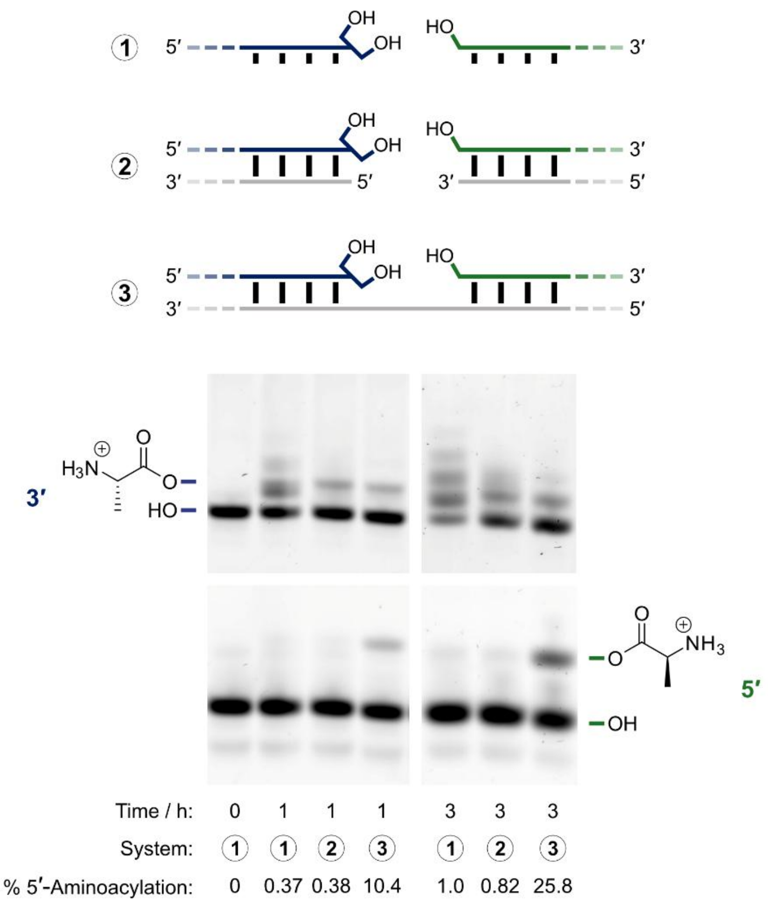


**Fig. S10 | Direct 5'-aminoacylation is not accessible from the thioester.** Thioester aminoacylations (MOPS (pH 8, 200 mM), NaCl (200 mM), Tween-20 (0.05%), L-Ala-SEt (200 mM)) were set up with three different oligonucleotide systems: *1* - FITCPriCCA (0.25 µM), dCy5PriA (0.25 µM); *2* - FITCPriCCA (0.25 µM), dCy5PriA (0.25 µM), TempA10 (0.275 µM), TempA5 (0.275 µM); *3* - FITCPriCCA (0.25 µM), dCy5PriA (0.25 µM), TempA (0.275 µM). Reactions were run in the same conditions as those used in the model system, at 20°C, pH 8, 200 mM thioester. dCy5PriA (the DNA analogue of Cy5PriA) was used in these reactions to deconvolute 5'-aminoacylation from internal 2'-aminoacylation. In system 1, terminal and internal aminoacylation is seen on the unbound RNA oligonucleotide FITCPriCCA and trace levels of direct 5'-aminoacylation are seen on the DNA oligonucleotide dCy5PriA. In system 2, binding to complementary RNA dampens internal aminoacylation on FITCPriCCA. In system 3, the transfer reaction is available, facilitating 5'-aminoacylation of dCy5PriA.

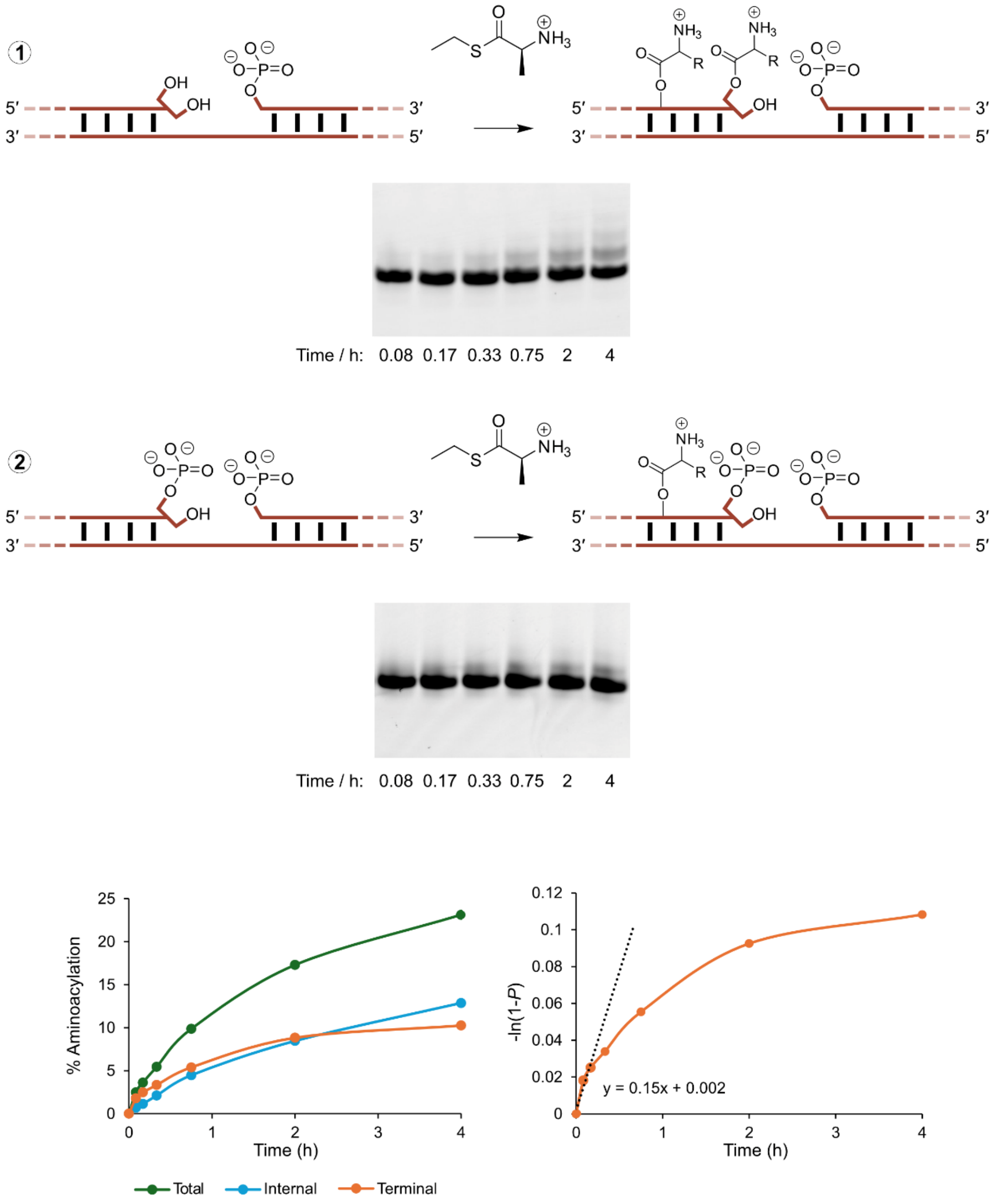


**Fig. S11 | Measuring the rate of thioester aminoacylation.** Thioester aminoacylation reactions (MOPS (pH 8, 200 mM), NaCl (200 mM), Tween-20 (0.05%), L-Ala-SEt (200 mM)) were set up with two oligonucleotide systems: 1 - FITCPriCCA (0.25 μM), pCy5PriA (0.25 μM) and TempA (0.275 μM); 2 - FITCPriCCAP (0.25 μM), pCy5PriA (0.25 μM) and TempA (0.275 μM). Both systems are incapable of transfer to the 5' in the junction; direct aminoacylation of the 2'/3'-diol in the junction (terminal aminoacylation; orange line, bottom left panel) is calculated by subtracting aminoacylation of the oligonucleotide with the 3'-phosphate (that blocks aminoacylation of the vicinal 2'-OH

(13)) in *2* (internal aminoacylation; light blue line, bottom left panel) from aminoacylation of the cis-diol version (total aminoacylation - including terminal and internal aminoacylation by thioester; green line, bottom left panel) in *1*. These estimates of 2'/3'-aminoacylation of FITCPriCCA were used to determine the initial rate. As the thioester aminoacylation rate decreases with time due to hydrolysis of the thioester, only the early timepoints (up to 10 mins) were used for its estimation. For the calculation of the thioester concentration needed to maintain 64% 2'/3'-aminoacylation in the absence of transfer, the model system was used to calculate the requisite aminoacylation rate (1.27 $h^{-1}$), and it was assumed that rate would increase with respect to thioester concentration in a linear manner. An equivalent 8.4-fold increase in rate would therefore require 1.7 M Ala-SEt.

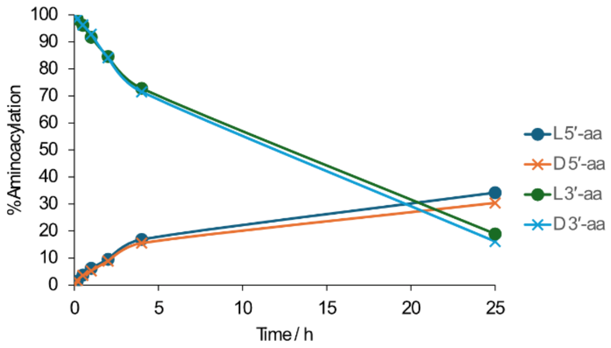


**Fig. S12 | The transfer reaction shows negligible stereoselectivity.** Plotted are L- vs D-Ala transfer reactions in MOPS (200 mM, pH 7), NaCl (200 mM), Tween-20 (0.05%), -7°C (frozen). Aminoacylation levels normalised to the starting level of 3'-L/D-Ala FITCPriCCA.

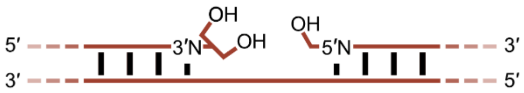


| 3′N | 5′N | % Transfer | $\Delta G^{\circ}_{37^{\circ}C}$* (kcal/mol) | $\Delta G^{\circ}_{37^{\circ}C}$† (kcal/mol) |
|---|---|---|---|---|
| A | A | 51.7 | -3.49 ± 0.14 | -0.93 ± 0.03 |
| U | A | 39.9 | -3.07 ± 0.13 | -1.33 ± 0.09 |
| A | U | 38.1 | -3.55 ± 0.14 | -1.10 ± 0.08 |
| G | A | 32.5 | -4.01 ± 0.17 | -2.35 ± 0.06 |
| A | G | 17.6 | -4.25 ± 0.38 | -2.08 ± 0.06 |
| A | C | 13.4 | -4.72 ± 0.58 | -2.24 ± 0.06 |
| C | A | 11.6 | -3.17 ± 0.16 | -2.11 ± 0.07 |

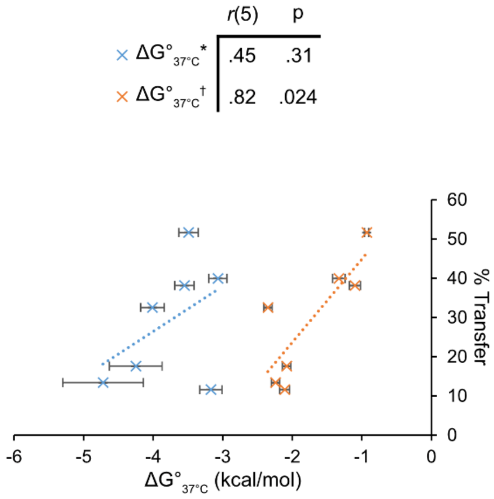


**Fig. S13 | The correlation between % transfer and two datasets of ΔG values of these sequences.** Pearsons's correlation coefficients were calculated between our measured % transfer values from fully Watson-Crick base paired nicks (in Fig. 3b, Fig. S8) and the same sequences' ΔG values for coaxial stacking free energies in a nick ('†', blue (14)) and nearest neighbour helix propagation free energies ( '*', orange (15)). Transfer extent is negatively correlated with both data sets, but significantly to the free energies used for nearest neighbour, helix propagation.

**Table S1 | Oligonucleotide Sequences**

DNA oligomers are shown in grey, RNA in black. For RNAs other than triplets generated by *in vitro* transcription, the two DNA primers shown were used to generate dsDNA by three rounds of mutual extension (GoTaq® Green Master Mix, Promega). Where multiple similar oligonucleotides differ only in the number of random nucleotides (N) contained at a specific sequence locus, these are represented by a single table entry. For these the sequence name is followed by a number range in square brackets e.g. NickT[0-5], each number in this range denotes an oligonucleotide containing that number of Ns. For instance, NickT3 contains NNN at the $[N]_{0-5}$ locus in the sequence. RNA triplets were generated by run-off transcription from a dsDNA with a 5' single-stranded overhang, formed by the two primers stated, as previously reported (16).

| Application | Name | Source | Sequence (5'→3') |
|---|---|---|---|
| PCR Primers | Primer 1 | Invitrogen | GATCGATCTCGCCCGCGAAATTAAT<br>ACGACTCACTATA |
| Flexizymes | eFx | Invitrogen | Forward: Primer 1<br>Reverse:<br>ACCTAACGCTAATCCCCTTTCGGG<br>GCCGCGGAAATCTTTCGATCCTATA<br>GTGAGTCGTATTAATTTCGCGGGCG<br>AGATCGATC<br>Transcript:<br>$^{PPP}$GGAUCGAAAGAUUUCCGCGGC<br>CCCGAAAGGGGAUUAGCGUUAGG<br>U |
| | dFx | Invitrogen | Forward: Primer 1<br>Reverse:<br>ACCTAACGCTAATCCCCTTTCGGG<br>GCCGCGGAAATCTTTCGATCCTATA<br>GTGAGTCGTATTAATTTCGCGGGCG<br>AGATCGATC<br>Transcript:<br>$^{PPP}$GGAUCGAAAGAUUUCCGCAUC<br>CCCGAAAGGGUACAUGGCGUUAG<br>GU |
| Donor Strands | FITCPriC<br>CA | IDT | FITC-UAGGAGACCA |

| | | | |
|---|---|---|---|
| | FITCpriCCC | IDT | FITC-UAGGAGACCC |
| | FITCpriCCG | IDT | FITC-UAGGAGACCG |
| | FITCpriCCU | IDT | FITC-UAGGAGACCU |
| | 3miHx | IDT | CAGAGCCGCCA |
| | FITCPriCCAP | IDT | FITC-UAGGAGACCA-3'P |
| Acceptor Strands | Cy5PriA | IDT | ACUGC-Cy5 |
| | dCy5PriA | IDT | ACUGC-Cy5 |
| | Cy5PriU | IDT | UCUGC-Cy5 |
| | Cy5PriC | IDT | CCUGC-Cy5 |
| | Cy5PriG | IDT | GCUGC-Cy5 |
| | FITC5miHx | IDT | GGCUCUG-FITC |
| | HO10Cy5 | IDT | GCGGGUGCAA-Cy5 |
| | pCy5PriA | IDT | $^{P}$ACUGC-Cy5 |
| Template Strands | TempA | IDT | GCAGUUGGUCUCCUA |
| | dTempA | IDT | GCAGUUGGUCUCCUA |
| | TempU | IDT | GCAGAUGGUCUCCUA |
| | TempC | IDT | GCAGGUGGUCUCCUA |
| | TempG | IDT | GCAGCUGGUCUCCUA |
| | TempCA | IDT | GCAGUGGGUCUCCUA |
| | TempGA | IDT | GCAGUCGGUCUCCUA |
| | TempUA | IDT | GCAGUAGGUCUCCUA |
| | TempCCACCC | Merck | GGGUGGUCUCCUA |
| | NickA[0-5] | Invitrogen | Forward: Primer 1<br>Reverse: CTTCGGAGTCTAGGAGACCA[N]$_{0-5}$ACTGCGCACCCGAAGGTGCTATAGTGAGTCGTATTAATTTCGCGGGCGAGATCGATC<br>Transcript: GCACCUUCGGGUGCGCAGU[N]$_{0-5}$UGGUCUCCUAGACUCCGAAG |
| | sTQT2 | IDT | ACCCGCUGGUCU |
| | TempA10 | IDT | UGGUCUCCUA |
| | TempA5 | IDT | GCAGU |

| | | | |
|---|---|---|---|
| Triplets | pppCCA | Invitrogen | Forward: Primer 1<br>Reverse:<br>TGGTATAGTGAGTCGTATTAATTTCGCGGGCGAGATCGATC<br>Transcript:<br>$^{PPP}$CCA |
| | pppCCC | Invitrogen | Forward: Primer 1<br>Reverse:<br>GGTATAGTGAGTCGTATTAATTTCGCGGGCGAGATCGATC<br>Transcript:<br>$^{PPP}$CCC |
| | pCCA | Eurogentec | $^{P}$CCA |
| Competitors | CompA | IDT | UAGGAGACCAACUGC |
| | Loop1CCA | Merck | GACUCUUCGGAGUCUAGGAGACCA |
| | CompNN | IDT | UAGGAGACCNNCUGC |
